# Length-independent telomere damage drives cardiomyocyte senescence

**DOI:** 10.1101/394809

**Authors:** Rhys Anderson, Anthony Lagnado, Damien Maggiorani, Anna Walaszczyk, Emily Dookun, James Chapman, Jodie Birch, Hanna Salmonowicz, Mikolaj Ogrodnik, Diana Jurk, Carole Proctor, Clara Correia-Melo, Stella Victorelli, Edward Fielder, Rolando Berlinguer-Palmini, Andrew Owens, Laura Greaves, Kathy L. Kolsky, Angelo Parini, Victorine Douin-Echinard, Nathan K. LeBrasseur, Helen M. Arthur, Simon Tual-Chalot, Marissa J. Schafer, Carolyn M Roos, Jordan Miller, Neil Robertson, Jelena Mann, Peter D. Adams, Tamara Tchkonia, James L Kirkland, Jeanne Mialet-Perez, Gavin D Richardson, João F. Passos

## Abstract

Ageing is the biggest risk factor for cardiovascular health and is associated with increased incidence of cardiovascular disease. Cellular senescence, a process driven in part by telomere shortening, has been implicated in age-related tissue dysfunction. Here, we address the question of how senescence is induced in rarely dividing/post-mitotic cardiomyocytes and investigate if clearance of senescent cells attenuates age related cardiac dysfunction. During ageing, human and murine cardiomyocytes acquire a senescent-like phenotype characterised by persistent DNA damage at telomere regions that can be driven by mitochondrial dysfunction, and crucially can occur independently of cell-division and telomere length. Length-independent telomere damage in cardiomyocytes activates the classical senescence-inducing pathways, p21^CIP^ and p16^INK4a^ and results in a non-canonical senescence-associated secretory phenotype. Pharmacological or genetic clearance of senescent cells in mice alleviates myocardial hypertrophy and fibrosis, detrimental features of cardiac ageing, and promotes cardiomyocyte regeneration. Our data describes a mechanism by which senescence can occur and contribute to ageing in post-mitotic tissues.

## Introduction

The role of cellular senescence in tissue maintenance, repair, and ageing is currently a rapidly evolving area of research under intense focus. Cellular senescence is classically defined as an irreversible loss of division potential of mitotic cells, often accompanied by a senescence-associated secretory phenotype (SASP). Whilst senescence can be detrimental during the ageing process, it has also been implicated in fundamental biological processes such as tumour suppression, embryonic development, and tissue repair (van Deursen, 2014).

Telomere shortening has been proposed as a major inducer of senescence (Bodnar, Ouellette et al., 1998). Telomeres are repetitive sequences of DNA, associated a protein complex known as shelterin (de Lange, 2005) that facilitates formation of a lariat-like structure to shield the exposed end of DNA (Griffith, Comeau et al., 1999), thus protecting the ends of chromosomes from being recognised as DNA damage (d’Adda di Fagagna, Reaper et al., 2003). The current dogma of telomere biology suggests that telomere shortening with each successive cell division eventually disrupts the protective cap, leading to a sustained DNA-damage response (DDR) and activation of the senescence programme (Griffith et al., 1999). This hypothesis may explain age-related degeneration of tissues maintained by constant contribution of stem cell pools, such as the skin and hematopoietic systems; however, it is insufficient to explain how senescence contributes to ageing in primarily post-mitotic tissues such as the heart. As such, the mechanisms that drive senescence in postmitotic cells and the contribution that postmitotic cell senescence (PoMiCS) in tissue degeneration, including the heart, during ageing is an emerging area of interest (Anderson, Richardson et al., 2018, Sapieha & Mallette, 2018).

Heart failure is a common disease of the elderly. Distinct physiological and molecular changes occur in the normal heart with ageing, including cardiomyocyte (CM) hypertrophy, which is associated with myocardial dysfunction and can progress to clinical heart failure (Strait & Lakatta, 2012). Critically short telomeres, induced by breeding of multiple generations ageing of mice lacking the catalytic subunit of telomerase *Terc*, leads to cardiac dysfunction and myocardial remodelling (Wong, Oeseburg et al., 2008). However, replicative senescence induced telomere shortening is unlikely to reflect normal physiological myocardial ageing, considering that the majority of CMs are postmitotic, withdrawing from the cell cycle shortly after birth. While the heart has a limited potential for regeneration, CM turnover rates in both humans and mice are extremely low (Bergmann, Zdunek et al., 2015, Richardson, Laval et al., 2015, Senyo, Steinhauser et al., 2013). We and others have reported that stress induced telomere damage can also lead to telomere dysfunction and induce senescence (Hewitt, Jurk et al., 2012). Therefore, we investigated if this phenomenon occurs in CMs, whether this provides a mechanism for PoMiCS and if CM senescence contributes to myocardial remodelling during ageing. Here, we demonstrate for the first time that a CM senescence-like phenotype is a feature of normal physiological human and murine ageing and provide a novel mechanistic model by which senescence occurs in rarely dividing/post-mitotic tissues. Persistent DNA damage foci, which co-localize with telomere regions, increase in cardiomyocytes with age independently of telomere length, telomerase activity, or DNA replication and can be induced by mitochondrial dysfunction *in vitro* and *in vivo*. Global gene expression analysis of purified CMs, isolated from young and old mice, indicates activation of classical senescence growth arrest pathways and aged CMs produce a distinct non-canonical SASP with functional effects. Furthermore, specific induction of length-independent telomere dysfunction in CMs induces several senescence markers, including hypertrophy, which importantly is also associated with CM senescence *in vivo*. Finally, we demonstrate that suicide gene-mediated ablation of p16^Ink4a^ expressing senescent cells and treatment with the senolytic drug, navitoclax, reduces CMs containing dysfunctional telomeres, promotes CM regeneration, attenuates cardiac hypertrophy and reduces fibrosis in aged mice.

## Results

### Length-independent telomere damage in aged cardiomyocytes *in vivo*

Previously, we defined a novel mechanism of cellular senescence *via* the induction of irreparable telomere damage that occurs in the absence of telomere shortening (Hewitt et al., 2012). These data led us to hypothesise that this form of telomere damage could be the process by which senescence occurs in post-mitotic, or rarely dividing cells, which are not subject to proliferation-associated telomere shortening. We therefore investigated telomere dysfunction in adult mice throughout ageing by using dual Immuno-FISH to quantify co-localisation between DDR proteins γH2A.X, 53BP1 and telomeres, hereafter referred to as Telomere-Associated Foci (TAF) in CMs (**Figure 1a**). CMs were identified by CM specific markers α-actinin, troponin C, and the perinuclear protein PCM1 (Richardson, 2016, Richardson et al., 2015) (**Fig S1a-c**).

**Figure 1.**
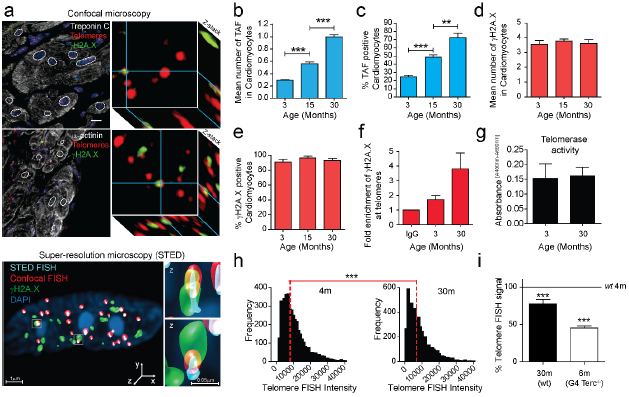
Telomere dysfunction increases with age in mouse cardiomyocytes. **(a)** Representative images of γH2AX immuno-FISH in troponin-C/α-actinin (top left and bottom left panels, respectively) positive mouse CMs (white – troponin-C/α-actinin; red – telo-FISH; green – γH2AX). Images are Z-projections of 0.1μM stacks taken with a 100x oil objective. Scale bar: 10μM. Right panels represent a single Z-plane where co-localisation between a γH2AX foci and telomere was observed. (below) 3D reconstruction of Immuno-FISH using STED microscopy for γH2A.X and telomeres in a CM from 30 month old mice. **(b)** Mean number of TAF and **c)** % of TAF positive α-actinin positive CMs from C57BL/6 mice. Data are mean ± SEM of n=4-7 mice per age group. 100 α-actinin positive CMs were quantified per mouse. **(d)** Mean number of total γH2A.X foci and **e)** % of γH2A.X foci-positive nuclei (bottom) in α-actinin positive CMs from C57BL/6 mice. Data are mean ± SEM of *n*=4-7 mice per age group. 100 α-actinin positive CMs were quantified per mouse. **(f)** Fold enrichment of γH2AX at telomere repeats by real-time PCR. Graph represents fold enrichment of γH2AX at telomeric repeats between IgG control, 3 and 30 month whole mouse hearts; Data are mean ± SEM of n=3 mice per age group. **(g)** Quantitative PCR-ELISA TRAP assay comparing telomerase activity of 3 and 30 month old C57BL/6 mice whole heart lysates. Data are mean ± SEM of n=4 mice per age group. **(h)** Histograms displaying distributions of telomere intensity in CMs analysed by 3D Q-FISH in young (4 months) and old (30 months) wild-type mice. Data are from n=3 mice. > 100 CMs were analysed per mouse. **(i)** % of telomere FISH signal loss in 30 months old wild-type mice and late generation Terc^-/-^ mice (6 months old) in comparison to 4 months old wild-type mice. Data are from n=3 mice. >100 CMs were analysed per mouse. Statistical analysis was performed using One Way ANOVA (Holm-Sidak method) for multiple comparisons and Mann-Whitney test and two-tailed t-test for single comparisons. ***P<0.0001; **P<0.01; * P<0.05.

Mean TAF per CM and the total % of CMs containing TAF increased significantly with age in both mouse and human hearts (**Figure 1a-c and Fig S1a**). In contrast, the total number of γH2A.X foci did not change with age and were detected in almost all adult CMs irrespective of age (**Figure 1d & e**). Similar results were obtained when analysing co-localisation between DDR protein 53BP1 and telomeres in young and old mice (**Fig S1b**). Interestingly, CMs from old animals contained a significantly higher number of TAF than other cardiac cell-types **(Fig S1c),** suggesting that telomere dysfunction is a predominant feature of CM ageing. To confirm further the localisation of a DDR at telomeres, we performed chromatin immunoprecipitation (ChIP) on cross-linked chromatin from heart tissue, using an antibody against γH2A.X followed by quantitative real-time PCR for telomeric repeats. We found enrichment of γH2A.X at telomeres in 30 month compared to 3 month old mice (**Fig 1f**).

Although the majority of the adult CMs are considered post-mitotic, the adult heart retains limited potential for CM proliferation (Bergmann et al., 2015, Richardson et al., 2015, Senyo et al., 2013). We next investigated if increased TAF was the outcome of telomere shortening driven by replication and/or impaired telomerase activity. Having first observed that telomerase activity was not affected by mouse age (**Figure 1g**), we observed a significant reduction in telomere FISH signal when comparing 4 month to 30 month old mice **(Figure 1h)**. However, the median telomere FISH signal detected in old wild-type (30 month) mice was comparatively higher than that found in 4^th^ generation Terc^-/-^ (G4) where critical telomere shortening has been shown to induce cardiac dysfunction (Wong et al., 2008) (**Figure 1h**).

3D super-resolution stimulated emission depletion microscopy (STED) was used to more accurately detect telomere length by Q-FISH in CMs. STED resolution is typically 50nm in XY and 150nm in Z and overcomes the so-called diffraction limit of conventional confocal microscopy which yields resolutions of ≥ 200nm. Using STED, we could detect an increased number of telomeres detected per CM, as well as decreased individual telomere volume when compared to conventional confocal microscopy (**Figure S1d & e and Extended Data Video**). STED microscopy also revealed the existence of clusters of telomeres with varying lengths, which would otherwise be identified as single telomere signals by confocal microscopy (**Figure S1d**).

We then proceeded to analyse telomere length by Q-FISH in CMs from old mice using STED. telomere FISH intensities in telomeres co-localising with γH2A.X (TAF) and those not co-localising with γH2A.X (non-TAF) in aged mice and found no significant differences **(Figure S1f & g)**. Similar observations were made in CMs from aged human hearts (**Figure S1h**). Analysis of telomere FISH intensities by STED in individual cardiomyocytes revealed that telomeres with low FISH intensity co-localising with γH2A.X are relatively rare **(Figure S1i)**. Together, our data suggest that length-independent telomere-associated DNA damage is not due to the limitations of conventional confocal microscopy and that the shortest telomeres are not preferentially signalling a DDR.

While CM proliferation is negligible in adult humans and mice, it is possible that TAF may be the outcome of a small fraction of dividing CMs. To ascertain further if age-associated accumulation of TAF, can occur in post-mitotic cells and is not exclusively linked to CM proliferation and turnover during ageing, we analysed our data in a mathematical model of a CM population over 27 months of a mouse lifespan. Our simulations reveal that, even considering relatively high estimates of cell division rates (4-15%), CM proliferation cannot account for the rate of the age-dependent increase in TAF. Rather, if we assume that proliferation plays no role in the TAF increase with age and that TAF are generated in non-dividing cells, there is a high degree of correlation between the model and our experimental data (**Figure S2**). In the *mdx*^*4cv*^*/mTR*^*G2*^ mouse model of telomere dysfunction, reduced expression of shelterin components is suggested to underlie increased telomere erosion in CMs (Chang, Ong et al., 2016, Mourkioti, Kustan et al., 2013). To test if uncapping of telomeres contributes to induction of a DDR at telomeres during ageing, we quantified the expression of shelterin components in CMs isolated from young and old wild-type mice and found no significant differences. (**Figure S3a**). Similarly, we observed no significant difference in the abundance of TRF2 (an essential component of t-loop formation) in either TAF or non-TAF in human CMs *in vivo* (**Figure S3b**).

Together this data supports the notion that TAF increase with age in CMs and this occurs as a result of a process that is independent of cell proliferation, can occur independently of telomere shortening and is not a result of overt alteration of telomere regulatory factors, such as shelterin components and telomerase. Having shown the phenomenon of telomere dysfunction occurring in CMs *in vivo*, we also found an age-dependent increase in TAF (but not non-TAF) was observed in other post-mitotic cells; specifically in skeletal muscle myocytes and hippocampal neurons which indicates the widespread nature of this phenomenon (**Figure S4**).

### Telomere damage is persistent in cardiomyocytes

Our previous work demonstrated that stress-exposure leads to activation of a persistent DDR at telomeric regions (Fumagalli, Rossiello et al., 2012, Hewitt et al., 2012). To investigate the kinetics and nature of the DDR in CMs, we utilised different *in vitro* models. We first identified that exposure to X-ray radiation (10Gy) resulted in both Telomere-Associated Foci (TAF) and non-Telomere Associated DNA damage Foci (non-TAF) in mouse embryonic CMs positive for troponin C and PCM1 **(Figure 2a)**. However, only TAF were persistent, with non-TAF numbers being significantly reduced with time **(Figure 2b)**.

**Figure 2.**
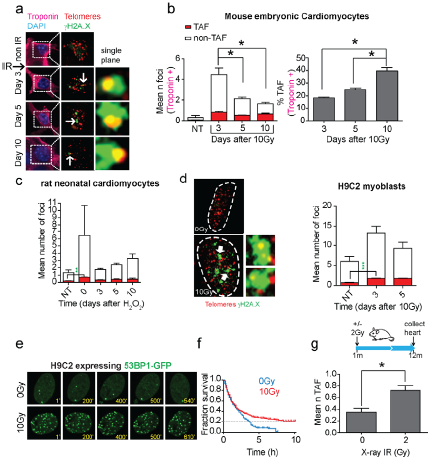
Stress-induced telomere-associated DNA damage is persistent in mouse embryonic cardiomyocytes, rat neonatal cardiomyocytes and H9C2 myoblasts. **(a)** Representative images of mouse embryonic cardiomyocytes at days 0, 3, 5 and 10 days following 10Gy X-irradiation. Left panels represent troponin C-positive embryonic cardiomyocytes (troponin C – magenta; DAPI - light blue). Middle panels display γH2AX foci (green) and telomeres (red) in Z projections of 0.1μM slices, with white arrows indicating co-localisation. Co-localising foci are amplified in the right-hand panels (amplified images represent a single Z-plane where co-localisation was observed). **(b)** (left) Mean number of both TAF and non-TAF in troponin I-positive mouse embryonic cardiomyocytes at days 0, 3, 5 and 10 following 10Gy X-irradiation. Data are mean ±S.E.M of *n*=3 independent experiments; 30-50 troponin positive cardiomyocytes were analysed per experiment. (right) Mean percentage of γH2AX foci co-localising with telomeres (% TAF) in troponin C-positive mouse embryonic cardiomyocytes at days 0, 3, 5 and 10 following 10Gy X-irradiation. Statistical analysis performed using One-Way ANOVA (Holm-Sidak method); * P<0.05. Significant differences were found for mean number of non TAF, but not for mean number of TAF. **c)** Mean number of both TAF and non-TAF (left graph) in neonatal rat cardiomyocytes at days 0, 3, 5, 10 days following treatment for 24h with H_2_O_2_. Data are mean ±S.E.M of *n*=3. >50 cells were quantified per condition; **d)** (left) Representative images of γH2AX immuno-FISH in H9C2 myoblasts 3 days 10Gy X-irradiation (red – telo-FISH; green – γH2AX) (right) Mean number of both TAF and non-TAF (left graph) in H9C2 myoblasts at days 3 and 5 following 10Gy X-irradiation. Data are mean ±S.E.M of *n*=3. >50 cells were quantified per condition; **e)** Representative time-lapse images of H9C2 rat cardiomyoblasts expressing AcGFP-53BP1 from 3 days after 10Gy irradiation at the indicated times (mins). Images are maximum intensity projections with a 6.7μM focal depth. **f)** Kaplan-Meyer survival curves for AcGFP-53BP1c DDR foci in H9C2 cells 3 days after 10Gy irradiation at 10 minute intervals for 24 hours. >500 foci from 10 cells were tracked per condition. Gehan-Breslow Test was used P<0.001. **g)** Schematic illustration showing 1 month old C57BL/6 mice were treated with 2Gy whole body X-irradiation, followed by a recovery period of 11 months before culling at 12 months of age. Mean number of TAF in α-actinin positive cardiomyocytes. Data are mean ± SEM of *n*=3 mice per group. 90 α-actinin positive cardiomyocytes were quantified per condition. Statistical analysis performed using two-tailed t test; * P<0.05

Cultures of rat neonatal CMs and the myoblast cell line (H9C2) treated with stressors H_2_O_2_ and X-ray irradiation, respectively, showed that the number of TAF remained persistent over time (**Figure 2c & d**). Three days subsequently to X-ray irradiation, we monitored DDR in H9C2 cells using a 53BP1-GFP fusion protein (Passos, Nelson et al., 2010) and time-lapse microscopy **(Figure 2e)**. Over a 10 hour time-course, the majority of individual DDR foci could be seen to be rapidly resolved with only approximately 20% of the original DNA damage foci persisting throughout the time course (**Figure 2f**), similar to the % of TAF positive cells for this time point (**Figure 2d**). To determine if persistent TAF can also be induced in adult CMs *in vivo*, we exposed young mice to a whole-body dose of 2Gy X-ray irradiation and measured the level of TAF in CMs after 11 months recovery. Both the mean number of TAF and % of CMs positive for TAF were significantly higher **(Figure 2g)**. Altogether, these data suggest that the majority of genomic DNA damage in CMs is reparable and only telomeric DNA damage is irreparable and persistent.

Double strand breaks can arise in telomeres due to replication errors when replication forks encounter single strand breaks. In order to ascertain if TAF could be induced in the absence of cell division, we pre-incubated H9C2 cells with EdU for 3 h and subjected them to either irradiation (10Gy) or 10μM H_2_O_2._ Following 24 h recovery (in the presence of Edu) the mean number of TAF was significantly increased in cells which did not incorporate Edu (**Figure S5a**), indicating that TAF can be induced in the absence of DNA replication. We further isolated adult mouse CMs, which do not proliferate in culture, and exposed them to 5μM H_2_O_2_. We observed that the mean number of TAF was significantly increased after treatment (**Figure S5b)**. Furthermore, we did not find evidence that the shortest telomeres were specifically targeted by stressors. In fact, EdU-negative irradiated H9C2 cells had a slightly higher median telomere length in damaged telomeres (TAF) compared to telomeres that did not co-localise with γH2A.X (non-TAF). However, such a difference was not found in adult murine CMs treated with H_2_O_2_ (**Figure S5c**).

### Telomere-specific DNA Damage Drives a Senescent-like Phenotype

Having demonstrated that TAF increase with age and can be induced due to stress, we then tested if telomere-specific damage could directly drive a senescent-like phenotype in the CM lineage. Rat neonatal CMs were transfected with an endonuclease that specially targets telomeres (a TRF1-FokI fusion protein (Dilley, Verma et al., 2016)). In culture, we found that rat neonatal CMs had very low proliferation rates with 5% of cells incorporating EdU over a period of 24h. Following 5 days of culture, CMs expressing TRF1-FokI showed numerous DNA damage foci with the significant majority of damage co-localising with telomeres (**Figure 3a, b & c**). Telomeres were targeted by the endonuclease irrespective of their length **(Figure 3d)**. *De novo* TAF formation induced a senescent phenotype in CMs characterised, in addition to TAF, by increased SA-β-Gal activity and upregulation of the cyclin-dependent kinase inhibitor p21^CIP^, as well as increased cellular hypertrophy (**Figure 3e-g**). Similar results were found using the H9C2 myoblasts (**Figure S6a-d**). Additionally, we used the AC10 cell line derived from adult human ventricular CM (Davidson, Nesti et al., 2005) stably expressing an inducible form of the TRF1-FokI construct. These cells express CM specific transcription factors and contractile proteins and, while proliferative in complete media, are induced to terminal differentiation and exit the cell cycle under mitogen-depleted conditions. We induced TRF1-FokI expression in differentiated AC10 cells by exposure to doxycycline (DOX) for 6h and allowed the cells recover for an additional 36h in the absence of DOX. Consistent with the idea that telomere damage is irreparable in differentiated CMs, we found that expression of TRF1-FokI significantly increased the mean number of TAF and this number persisted for at least 36 h (**Figure S6f & g**).

**Figure 3.**
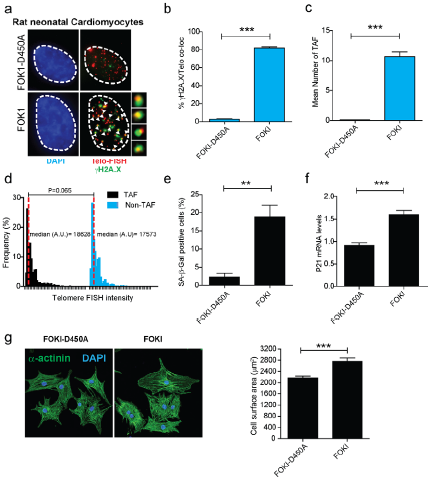
TRF1-FokI fusion protein induces telomere-specific double strand breaks, senescence, and hypertrophy in rat neonatal cardiomyocytes. **a)** Representative images of rat neonatal CMs 4 days following transfection with a FLAG-tagged TRF1-FokI-D450A (top row) or TRF1-FokI (middle and bottom row) fusion protein (cell treatments the same for all subsequent panels in Figure) (red – telo-FISH; green – γH2A.X). Images are z-projections of 0.1μM stacks taken with 100X objective. White arrows indicate co-localisation between telomeres and γH2A.X, with co-localising foci amplified in the right panels (taken from single z-planes where co-localisation was found).**b)** % of γH2A.X foci co-localising with telomeres and **c)** Mean number of Telomere-associated Foci (TAF) in FLAG-tagged TRF1-FokI-D450A- and TRF1-FokI-expressing CMs. Data are mean ± SEM of n=4 independent experiments. >50 cells were analysed per condition. Statistical analysis was performed by two-tailed t-test * P<0.05. **d)** Histograms displaying telomere intensity for telomeres co-localising or not co-localising with γH2AX foci. Red dotted lines represent median. Mann-Whitney test show no significant difference in telomere intensity between TAF and non-TAF. **e)** Mean % of FLAG-labelled CMs positive for SA-β-Gal activity. Data are mean ± SEM of n=3 independent experiments. >100 cells were quantified per condition. **f)** Expression of p21 mRNA (as a function of β-actin and Gapdh) by Real-time PCR in TRF1-FokI-D450A and TRF1-FokI expressing CMs. Data are mean ± SEM of n=6 independent experiments; **g)** Mean cell surface area (μm^2^) of FLAG-labelled CMs expressingTRF1-FokI-D450A and TRF1-FokI. Statistical analysis performed using two tailed t test; ***P<0.0001; **P<0.01.

### Aged cardiomyocytes activate senescent pathways but not a typical SASP

Within the heart, CMs comprise only approximately 30% of the total cells (Nag & Zak, 1979). To overcome this heterogeneity and obtain an accurate representation of the CM transcriptome during ageing, we devised a new method for isolation and enrichment of CMs **(Figure 4a**) to obtain a highly enriched CM population (**Figure S7a**). qRT-PCR quantification of mRNAs encoding the cyclin-dependent kinase inhibitors p16^Ink4a^, p21^CIP^, and p15^Ink4b^ in 3 and 20 month old animals demonstrated an age-dependent increase in expression of all three genes (**Figure 4b**). Immunohistochemistry on tissue sections from ageing mice validated the increase of p21^CIP^ at the protein level, specifically in CMs (**Figure 4c**). Furthermore, we detected increased activity of SA-β-Gal in old mice (**Figure 4d**). While SA-β-Gal positivity was rare, we could detect it in CMs but no other cell types from old mice. By centromere FISH in CMs, we also observed an age-dependent increase of Senescence-Associated Distension of Satellites (SADS), a marker of senescence (Swanson, Manning et al., 2013) (**Figure 4e**). Global transcription analysis of purified CM populations by RNA-sequencing revealed 416 differentially expressed genes between young (3 months) and old (20 months) (**Figure S7b**). Principal component analysis revealed that both cohorts separate well with the first and second components, accounting for 80.1% and 7.8% of the cumulative variance across the data-set (**Figure S7c**). Consistent with rare CM proliferation *in vivo*, we did not observe any significant changes in the expression of proliferation genes (**Figure S7d**). However, genes involved in the regulation of myofilaments, contraction, and CM hypertrophy were enriched in old mice (Myh7, Capn3, Ankrd1, Myom3, Myot, Mybpc2). Confirmatory qRT-PCR showed age-dependent increase in hypertrophy genes Myh7 and Acta1, but not Anf (**Figure S7e and f**). Interestingly, pro-inflammatory genes associated with the SASP (Coppé, Patil et al., 2008), such as Il1a, Il1b, Il6, Cxcl1, and Cxcl2, were not differentially expressed between CMs from young and aged animals (**Figure 4f**). To determine if SASP factors were secreted by CMs, we collected conditioned medium from isolated CMs from young and old mice. In accordance with the RNA-seq results, we did not find any significant differences in the levels of secreted proteins using a cytokine array (**Figure 4f**).

**Figure 4.**
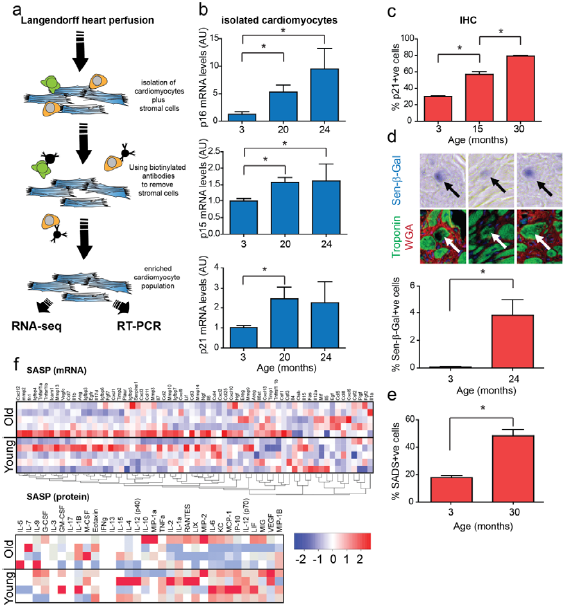
Aged cardiomyocytes activate senescent pathways but not a typical SASP. **a)** Schematic illustrating CM isolation procedure. **b)** Real-time PCR gene expression analysis in isolated mouse CMs from C57BL/6 mice. Data are mean ± SEM of *n*=3-4 per age group. Statistical analysis performed by One-Way ANOVA (Holm-Sidak method); * P<0.05. **c)** Mean % of p21-positive CM nuclei from 3, 15, and 30 month old C57BL/6 mice by Immunohistochemistry (IHC). Data are mean ± SEM of *n*=4 per age group. >100 CMs were quantified per age group. Statistical analysis performed using One-Way ANOVA (Holm-Sidak method); * P<0.05. **d)** Mean % of 3 and 24 month old mouse CMs staining positive for SA-β-Gal *in vivo* with representative images above (blue – SA-β-Gal; green – Troponin C; red – WGA). Statistical analysis performed using two-tailed t-test * P<0.05. Data obtained from the analysis of >500 CMs per mouse, 4 mice per age group. **e)** Mean % of SADS-positive CM nuclei from 3 and 30 month old mouse CMs positive for SADS *in vivo*, as detected by centromere-FISH. Data are mean ± SEM of *n*=4 per age group. >200 CMs per mouse were quantified. Statistical analysis performed using two-tailed t-test * P<0.05. **f)** SASP Heatmap: Pearson correlation clustered heatmap showing a curated list of known SASP genes (top panel) or a selection of secreted SASP proteins (bottom panel) in young (3 months) and old (20 months) mouse CMs (n=5 per age group). The colour intensity represents column Z-score, where red indicates highly and blue lowly expressed.

Our data are in contrast with a previous report that shows increased expression of SASP components in murine heart with age using whole heart homogenates (Ock, Lee et al., 2016). We therefore compared the expression of previously reported SASP components in non-purified and purified CM populations. Consistently, we found that following a simple *Langendorff* digestion that collect a heterogeneous population of CMs and stromal cells, we found significant differences in expression of SASP components such as Il-6 and Cxcl1 between young and old mice (**Figure S8a**). However, the population of purified CMs demonstrated no such differences, suggesting that cell-types other than CMs could explain previous observations (**Fig. S8a**). Interestingly, RNA sequencing led to identification of 3 secreted proteins, not commonly categorized as SASP components, which were confirmed to be significantly increased at the mRNA level in aged purified CMs: Edn3, Tgfb2, and Gdf15 (**Figure 5a**). Of these, only Edn3 was increased exclusively in aged CMs, but neither in cardiac stromal cells nor in other organs, suggesting that increased Edn3 levels in plasma is a good indicator of CM ageing (**Figure S8b-d**). The SASP has been shown to impact on proliferation of neighbouring cells, ultimately inducing senescence (Acosta, Banito et al., 2013). Consistently, we found that conditioned medium isolated from old adult CMs reduced proliferation of neonatal fibroblasts (measured by EdU incorporation) and increased the expression of α-smooth muscle actin (α-SMA), an indicator of myofibroblast activation (**Fig. 5b-e**). We then examined in more detail the role of the newly identified CM senescence-associated proteins in terms of their ability to induce bystander effects, particularly their role in fibrosis, proliferation, and hypertrophy. We found that Edn3, Tgfb2, and Gdf15 induced expression of α-SMA in fibroblasts, however only Tgfb2 reduced cellular proliferation as measured by EdU incorporation (**Figure 5f-h)**. Furthermore, treatment of neonatal rat CMs with Edn3 or Tgfb2, but not Gdf15, resulted in a significant increase in the cell surface area, supporting their involvement in the induction of hypertrophy (**Figure 5i**).

**Figure 5.**
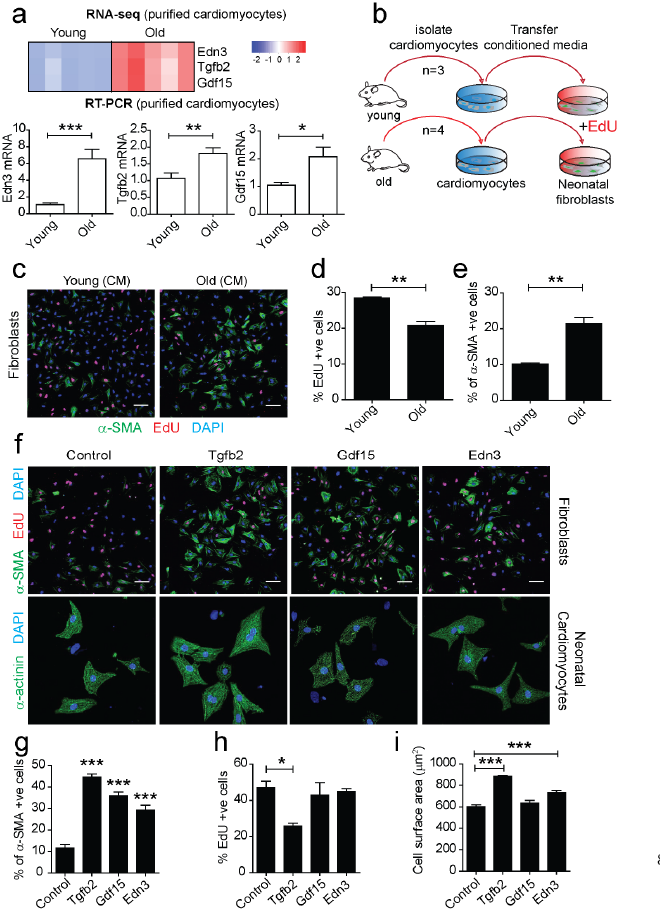
Cardiomyocyte SASP induces fibrosis and reduces proliferation in fibroblasts and induces hypertrophy in cardiomyocytes. **a)** (above) RNA-sequencing of purified CMs from 4 mice per age group reveal age-dependent increased expression of 3 secreted proteins Edn3, Tgfb2, and Gdf15; (below) mRNA expression of Edn3, Tgfb2, and Gdf15 was independently validated by RT-PCR in young and old isolated adult CMs. Data are mean ±S.E.M of n=8 mice per group. **b)** CMs were isolated by *Langendorff* heart perfusion, purified and cultured for 48h. Conditioned medium (CM) was collected and added to cultures of neonatal fibroblasts in the presence of 10μM EdU; **c)** Representative images of immunofluorescent staining against α-SMA and EdU in neonatal fibroblasts cultured in the presence of CM from young and old CMs; **d)** Quantification of % of EdU incorporation and **e)** % of a-SMA positive cells in neonatal fibroblasts after treatment for 48h with CM from young and old CM. Data are mean ±S.E.M of n=3-4 mice per age group; >200 cells were quantified per condition; **f)** Representative micrographs of neonatal fibroblasts and CMs treated with recombinant proteins: Tgfb2, Edn3, and Gdf15 for 48h and immunostained against α-SMA and EdU (fibroblasts) and a-actinin (CMs); **g)** Quantification of α-SMA and **h)** EdU positive neonatal fibroblasts and **i)** Surface area (μm^2^) in neonatal CMs following treatment with the indicated recombinant proteins. Data are mean ±S.E.M of n=3-4 independent experiments. Asterisks denote statistical significant with One-way ANOVA (multiple comparisons) or two-tailed t-test. ***P<0.0001; **P<0.01; * P<0.05.

We conclude that aged CMs *in vivo* activate a number of senescent effector pathways, including hypertrophy and a non-typical SASP, which may mediate both autocrine and paracrine effects.

### Mitochondrial ROS drives TAF in mouse cardiomyocytes *in vivo*

Mitochondrial dysfunction has been described as both a driver and consequence of cellular senescence (Correia-Melo, Marques et al., 2016, Passos et al., 2010, Passos, Saretzki et al., 2007b). Gene Set Enrichment Analysis (GSEA) of RNA-seq data revealed that the most negatively enriched Gene ontology term in old CMs is the mitochondrial inner membrane (**Figure S9a**). Consistently, in CMs from old mice we observed an overall decline in expression of most mitochondrial genes, particularly those genes involved in the Electron Transport Chain (ETC) (**Figure 6a**) and mitochondrial ultrastructural defects by transmission electron microscopy (**Figure S9b**).

**Figure 6.**
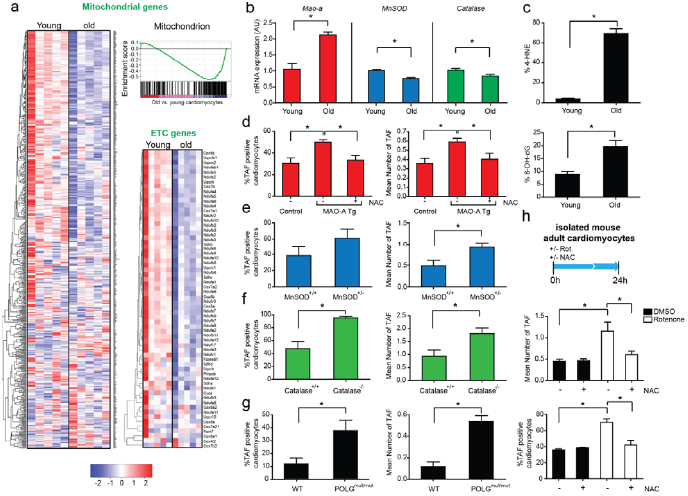
Mitochondrial dysfunction is a feature of cardiomyocyte senescence and drives TAF in mouse cardiomyocytes *in vivo*. **a)** Mito and ETC genes with GSEA analysis: Clustered heatmap showing all genes associated with the “Mitochondrion” GO term in young and old, mouse CMs as observed by the GSEA pre-ranked list enrichment analysis (normalised enrichment score: −1.70; FDR q-value < 0.05). Alongside this is a column clustered heatmap displaying a list of genes from the electron transport chain (ETC) GO ontology. In both instances, genes are by column and samples by row with the colour intensity representing column Z-score, where red indicates highly and blue lowly expressed. **b)** Real-time PCR gene expression analysis of MAO-A, MnSOD and catalase in isolated mouse CMs from young (3 months) and old (20 months) mice. Data are mean ± SEM of n=4-5 per age group. Asterisks denote a statistical significance at P<0.05 using two-tailed t-test. **c)** Mean % of 4-HNE-(top) or 8oxodG-(bottom) positive CMs from 3 month (young) or 30 month (old) aged mice. Data are mean ± SEM of n=4 per age group. 100 CMs were quantified per age group. Asterisks denote a statistical significance at P<0.05 using two-tailed t-test. **d-g)** Mean % of TAF-positive nuclei (left graphs) or mean % of TAF (right graphs) in wild-type (control) compared to MAO-A transgenic mice with or without drinking water supplemented with 1.5g/kg/day NAC from the age of 4 to 24 weeks, MnSOD^+/+^ vs MnSOD^-/+^, Catalase^+/+^ *vs*. Catalase^-/-^, WT *vs*. POLG double mutant mice. Data are mean ±S.E.M of n=3-4 per group. >100 CMs were quantified per age group. Statistical analysis was performed using two-tailed t test or One-way ANOVA (for multiple comparisons); * P<0.05. **h)** Schematic depicting isolated mouse adult CMs isolated from 4 animals were treated with or without 100nm rotenone either in the presence of 5mM NAC or vehicle control (pre-treated for 30 minutes before rotenone treatment), for 24 h before fixation. Mean number of TAF (top graph) and mean % of TAF-positive nuclei (bottom graph) from 4 separate CM cultures isolated from 3 month-old mice. 50 CMs were quantified per condition. Asterisks denote a statistical significance at P<0.05 using One-way ANOVA (Holm-Sidak method).

We then speculated whether mitochondrial ROS could be a driver of telomere dysfunction in CMs *in vivo*. Previous data indicate that mitochondrial ROS can drive telomere shortening *in vitro* (Passos et al., 2007b) and that telomere regions are particularly sensitive to oxidative damage (Hewitt et al., 2012). Consistent with this hypothesis, we found increased mRNA expression of the pro-oxidant enzyme monoamine oxidase A (MAO-A) and decreased expression of antioxidant enzymes mitochondrial MnSOD and catalase in isolated CMs from old mice (**Figure 6b**). Furthermore, we found increased lipid peroxidation marker, 4-HNE, and DNA oxidation marker, 8-oxodG, in hearts with age (**Figure 6c**). To address whether these age-associated changes are causal in TAF induction, we utilized a model of CM-specific overexpression of MAO-A (MHC-MAO-A tg), which displayed enhanced levels of mitochondrial ROS specifically in CMs (Villeneuve, Guilbeau-Frugier et al., 2013). In this transgenic mouse, both the mean number and % of CMs positive for TAF were significantly increased compared to age-matched controls (**Figure 6d**). Critically, when transgenic animals were treated with the antioxidant N-acetyl cysteine (NAC), both the increase in TAF (**Figure 6d**) and CM hypertrophy were significantly rescued (**Figure S9c**). Complementary studies in old MnSOD^+/-^ and Catalase^-/-^ mice also revealed that these had a higher number of TAF than aged-matched controls (**Figure 6e & f**). Furthermore, we found a significant increase in TAF in CMs from Polg^mut/mut^ mice, a model of accelerated ageing due to mitochondrial dysfunction (Trifunovic, Wredenberg et al., 2004) (**Figure 6g**). Increased telomere dysfunction was associated with increased expression of p21, cardiac hypertrophy, and decreased expression of mitochondrial proteins (**Figure S9d**). Finally, treatment of isolated adult CMs with rotenone, a mitochondrial complex I inhibitor, induced TAF, which could be rescued by treatment with the antioxidant NAC (**Figure 6h**).

In summary, CMs from aged heart exhibit down regulation of mitochondrial inner membrane and electron transport genes, and this is associated with indicators of increased ROS metabolism that is causative of telomere-associated DNA damage.

### Senescent-cell clearance reduces cardiac hypertrophy and fibrosis

Cardiac hypertrophy occurs with ageing. Furthermore, it accompanies diseases such as ischemia, hypertension, valvular disease, and heart failure. In order to assay cardiac function with ageing in mice we used Magnetic Resonance Imageing (MRI). We found no significant difference in cardiac function (ejection fraction). However, we observed an increased mean left ventricle mass (mean of diastolic and systolic mass) indicative of hypertrophy and increased ventricle wall rigidity, symptomatic of a decline in diastolic dysfunction, both of which are documented characteristics of cardiac ageing in humans and mice (Dai, Chen et al., 2012) (**Fig. 7a**).

**Figure 7.**
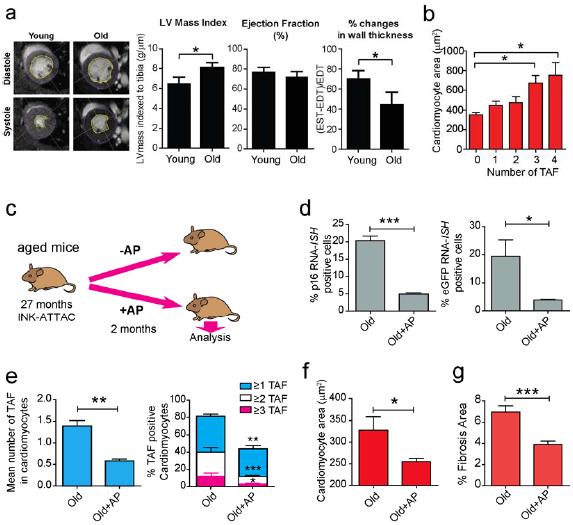
Genetic clearance of p16 positive cells in aged mice reduces cardiomyocyte senescence, cardiac hypertrophy and fibrosis. **a)** Examples of individual short axis cine-MR images of 3 or 20 month mouse hearts. Ejection fraction and LV thickness was calculated based on manual measurements of Left ventricle epicardial and endocardial borders. % change in wall thickening calculation based on wall thickness at the 4 points indicated. Measurements were made in all cine slices at end diastole and end systole. Graphs representing data obtained from MRI analysis of >7 animals per age group. Data are mean ± S.E.M. **b)** Comparison between mean number of TAF per CM and CM area in 22 month old animals. Data are mean ± S.E.M. of n=4. >100 CMs were quantified per mouse. Asterisks denote a statistical significance at P<0.05 using one-way ANOVA. **c)** Schematic depicting experimental design for Figures d-f: 27 month old INK-ATTAC mice were treated 4 times with AP20187 (or Vehicle), 3 days in a row, every 2 weeks (2m-long treatment in total) and were sacrificed afterwards for analysis. **d)** Comparison between the % of p16- or –eGFP positive CMs by RNA-*in situ hybridization* per plane in INK-ATTAC mice (28-29m old) treated with vehicle or AP20187. Data are mean ± SEM of *n*=5 per age group. 100 CMs were analysed per mouse. Asterisks denote a statistical significance at P < 0.05 using two-tailed t-test. **e)** Mean number of TAF (left graph) and mean % of TAF-positive nuclei (right graph) in CMs. Data are mean ± S.E.M. of n=6 per age group. 100 CMs were analysed per mouse. **f)** Histograms showing distribution of individual telomere intensities measured by Q-FISH in INK-ATTAC mice (28-29m old) treated with vehicle or AP20187. >150 CMs were analysed per mouse. **g)** Mean CM area μm^2^. Data are mean ±S.E.M. of *n*=6 per age group, >150 CMs analysed per mouse. **h)** % of fibrotic area evaluated by Sirius Red staining. Data are mean ±S.E.M. of n=6 per age group. Asterisks denote a statistical significance using two-tailed t test or Mann Whitney test. ***P<0.0001; **P<0.01; * P<0.05.

Previously, we showed that specific induction of TAF was associated with increased cell size in rat neonatal CMs *in vitro* and that CM senescence is associated with hypertrophy *in vivo*. Consistent with a role for TAF-induced senescence in age-dependent cardiac hypertrophy, we found that larger CMs from old mice had generally higher TAF frequencies (**Fig. 7b**). In order to determine if there is a causal relationship between senescence and cardiac hypertrophy, we used the INK-ATTAC mouse model, in which a small molecule, AP20187 (AP), induces apoptosis through dimerization of FKBP-fused Casp8. Using this model, it has previously been shown that clearance of p16-expressing cells improves multiple parameters of physical health and function within ageing mice (Baker, Childs et al., 2016, Baker, Wijshake et al., 2011, Farr, Xu et al., 2017, Ogrodnik, Miwa et al., 2017).

In order to establish if elimination of p16^Ink4a^ positive cells reduced TAF in CMs, we aged INK-ATTAC mice until they were 27 months old and treated them with AP for 2 months (**Figure 7c**). We confirmed by RNA-*in situ hybridization* that both p16 and eGFP positive cells were significantly reduced in hearts following AP treatment (**Fig. 7d**). Additionally, we found that TAF in CMs were significantly reduced (**Fig. 7e**), however, we failed to detect any differences in telomere FISH signals between INK-ATTAC mice with or without clearance of p16^Ink4a^-positive cells (**Figure S11a**). Consistent with a role for CM senescence in age-dependent myocardial remodeling, we found that AP treatment significantly reduced the average cross-sectional area of CMs (**Figure 7f**), and decreased the percentage area of fibrosis, without any change in heart function as measured by % ejection fraction (**Figure 7g and Figure S10b & c**). Similar results were observed in a model of cardiac hypertrophy induced by thoracic irradiation. We found that TAF induced by thoracic irradiation in INK-ATTAC mice were restored to the levels found in sham irradiated mice following AP treatment (**Figure S11a-c**). Similarly, we found that irradiation resulted in CM hypertrophy and this was completely rescued by elimination of p16 ^ink4a^ positive cells (**Fig. S11d**).

To investigate further the therapeutic impact of targeting senescent cells to counteract cardiac ageing, we treated aged wild-type mice with the previously described senolytic drug, ABT263 (navitoclax) (Zhu, Tchkonia et al., 2016) or vehicle intermittently for 2 weeks (**Figure 8a**). We found that navitoclax reduced telomere dysfunction in cardiomyocytes without affecting telomere length (similarly to what was observed in aged INK-ATTAC mice following AP treatment) **(Figure 8b, c and Figure S12a**). Consistent with its senolytic properties, we found that navitoclax selectively killed senescent H9C2 cardiomyoblasts *in vitro* (**Figure S12 b & c**).

**Figure 8.**
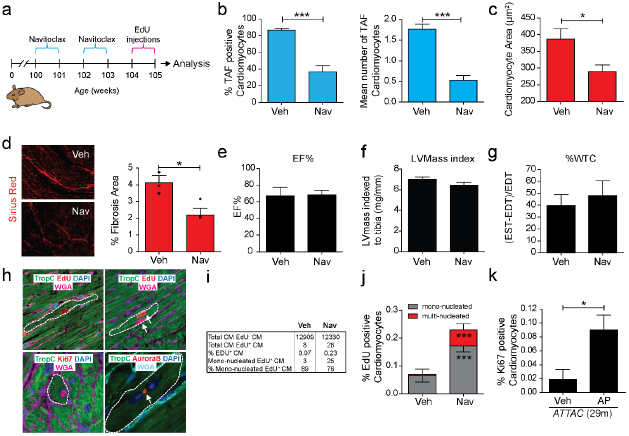
Pharmacological clearance of senescent cells with Navitoclax (ABT 263) reduces cardiomyocyte senescence and stimulates cardiomyocyte regeneration. **(a)** Schematic depicting experimental design. Mice at 100 weeks (23-months) of age were treated with vehicle (Veh) or navitoclax (Nav) intermittently for 2 weeks. At 104 weeks mice were injected every day with EdU for 1 week.; **b)** Quantification of mean number of TAF and % of TAF-positive CMs in 24 months old wild-type mice treated with vehicle or navitoclax (50mg/kg/day). Data are mean± SEM of *n*=5-8 mice per group. More than 100 CMs were quantified per animal; **c)** CM cross-sectional area in 24 months old wild-type mice treated with vehicle or navitoclax. Data are mean± SEM of *n*=8 mice per group; **d)** (left) representative images of Sirius red staining and (right) % of fibrosis area in 24 month-old mice treated or not with navitoclax. Data are mean mean±S.E.M of n=3 per group. **(e-f)** MRI analysis of Ejection Fraction (EF%), Left ventricle mass (LVmass) index and % of ventricle wall thickness (%WTC) in 24 month-old mice treated or not with navitoclax. Data are mean±S.E.M of n=6 mice per treatment group. **h)** Examples of confocal microscopy images of CMs positive for CM marker troponin C (TropC), EdU, Ki67 and Aurora B from navitoclax treated animals; **f)** Table summarising numbers of CMs quantified in j); **j)** Quantification of EdU positive CMs (mono- or multi-nucleated) in vehicle and navitoclax treated animals. Data are mean±S.E.M of n=5-6 mice per group; **k)** % of Ki67 positive CMs in 29 month old INK-ATTAC mice treated with vehicle or AP20187. Data are mean±S.E.M of n=4-6 mice per group. Asterisks denote a statistical significance using two-tailed t test or Mann Whitney test. ***P<0.0001; **P<0.01; * P<0.05.

Similarly to genetic clearance of p16^Ink4a^ cells in ATTAC mice, we found that navitoclax significant reduced hypertrophy and fibrosis in aged wild-type mice (**Figure 8c & d**). However, navitoclax had no significant impact on cardiac function, LV mass and ventricle wall rigidity (**Figure 8e-g**). The decrease in mean CM size without significant changes in LV mass suggested a compensatory increase in overall CM number. Supporting *de novo* CM proliferation, we observed that frequency distribution analyses of CM cross-sectional area suggested that the decrease in mean CM area following navitoclax treatment, is a function of both an elimination of the largest CMs, presumably as these are senescent, as well as the appearance of a “new” population of small CMs (**Fig. S12e**). This phenotype has previously been associated with CM regeneration (Waring, Vicinanza et al., 2014). To further investigate *de novo* CM regeneration following navitoclax treatment, we injected EdU in 24 month old mice for 7 days and quantified EdU positive cells in combination with CM markers TropC and PCM1. We found significantly more EdU labelled CMs in the Navitoclax treated animals than vehicle controls (0.23% vs 0.07%) (**Figure 8h-j**). As CMs may undergo karyokinesis in the absence of cytokinesis resulting in bi- or multi-nucleation, EdU alone is not a marker of cell division. As such, we evaluated the nucleation state of EdU labelled CMs by analyzing 3D images in 40μm thick sections as previously described (Mollova, Bersell et al., 2013). Our data indicates senescent cell clearance is accompanied by a significant increase of both multi- and mono-nucleated CMs (**Figure 8h-j**). However, less than 25% of all EdU incorporation resulted in multi-nucleated CMs. We next evaluated the % of CMs positive for proliferation marker Ki67 and found a similar increase following navitoclax treatment and in aged INK-ATTAC mice treated with AP (**Figure S12d and Figure 8k**). Expression of Aurora B, a component of the contractile ring required for cytoplasmic separation during cell division, was only observed in CMs from hearts where senescent cells had been cleared (**Figure 8h**). Altogether our data support that senescent CMs are involved in age-associated cardiac hypertrophy and fibrosis and that their clearance may induce CM regeneration.

## Discussion

During ageing and despite lows level of proliferation, we observe enrichment of DDR proteins at telomere regions, notoriously known for their inefficient repair capacity, fuelled by the actions of telomere binding proteins. TRF2 and its binding partner RAP1 have been shown to prevent NHEJ-dependent telomeric DNA fusions by inhibiting DNA-PK and ligase IV mediated end-joining (Bae & Baumann, 2007). In contrast, homologous recombination can repair telomere-induced DNA damage, however this process is restricted to proliferating cells undergoing S-phase and therefore not likely to be relevant for post-mitotic CMs (Mao, Liu et al., 2016). As such, within CMs, a cocktail of age-dependent mitochondrial dysfunction and oxidative stress coupled with limited CM turnover, fuels the occurrence of irreparable telomere-associated damage, which may instigate a senescent-like phenotype (**Figure S13**).

While our data indicate that mitochondrial dysfunction and ROS can induce CM senescence, it remains possible that the observed age-dependent changes in mitochondrial gene expression and morphology are a consequence of telomere dysfunction. In mouse models of accelerated telomere shortening, critically short telomeres have been shown to repress PGC1-α and β mediated mitochondrial biogenesis (Chang et al., 2016, Sahin, Colla et al., 2011). However, increased mitochondrial ROS driven by telomere dysfunction has been shown to induce DNA damage and activate a DDR in a positive feedback loop, making it experimentally complex to unravel which is the initiating step in the process (Passos et al., 2010).

There is still uncertainty regarding the physiological role of post-mitotic CM senescence during the ageing process. Previously studies have attributed the beneficial effects of clearance of senescent cells in murine ageing heart to clearance of senescent cells in proliferation-competent cells and not CMs (Baker et al., 2016). Studies have also proposed a role for telomere shortening in CM senescence: short telomeres have been observed in aged murine CMs (Rota, Hosoda et al., 2007) and mouse models of accelerated telomere shortening have cardiac dysfunction (Chang et al., 2016). However, whether short telomeres *per se* are causal in cardiomyocyte senescence during natural ageing has not been determined. Our data suggest that telomere length is not a limiting factor in CM senescence since: i) genetic elimination of p16^Ink4a^ positive cells or treatment with senolytics, navitoclax in aged mice did not affect telomere length or the frequency of very short telomeres; ii), super-resolution STED microscopy, that effectively resolved clustered telomere signals, could not detect differences in FISH intensity between telomeres co-localizing or not with γH2A.X. However, we acknowledge that there are limitations to our experimental approach: while interphase telomere FISH allows us to collect information regarding telomere length in specific cells of interest, it does not allow the recognition of telomeres in specific chromosomes or the detection of telomere-free ends.

Key questions remain regarding the causes and consequences of CM senescence. While telomere dysfunction in CMs activates the classical senescence-inducing pathways, p21^CIP^ and p16^Ink4a^, CMs in contrast to cardiac stromal cells they do not produce a classical SASP. Rather we have identified that factors such as Edn3, Tgfb2, and Gdf15 are released by senescent CMs and promote myofibroblast activation, contribute to CM hypertrophy and may constitute part of a CM specific SASP.

In summary, our study provides evidence for the concept that post-mitotic CM senescence is a major effector of myocardial ageing and offers a proof-of-principle that modulation of cardiac senescence is a viable treatment strategy. While described in myocytes, we speculate that our proposed mechanism may explain PoMiCS which has been detected in other tissues such as neurones (Jurk, Wang et al., 2012), osteocytes (Farr et al., 2017), and adipocytes (Minamino, Orimo et al., 2009).

## Material and methods

### Animals and Procedures

#### Ageing studies

Mixed sex C57BL/6 mice were analysed at either 3, 15, 20, 24, or 30 months of age. For whole body X-ray irradiation, mixed sex 1 month old C57BL/6 mice were irradiated with 2Gy followed by 11 month recovery period before culling. *TERC*^*-/-*^ *C57BL/6 mice.* Male mice were bred to produce successive generations of mice with decreasing telomere length. Hearts from fourth-generation (G4) mixed sex, mice were studied. *Catalase*^*-/-*^ *and MnSOD*^*+/-*^ *Mice*. Mouse models for elevated reactive oxygen species and mitochondrial dysfunction: Catalase^-/-^ and MnSOD^+/-^ were aged until 22-24 months. *PolgA*^*mut/mut*^ *mice*. Mice with a knock-in missense mutation (D257A) at the second endonuclease-proofreading domain of the catalytic subunit of the mitochondrial DNA (mtDNA) polymerase Polγ (*PolgA*^*mut/mut*^ mice) and *PolgA*^*+/+*^ mice were used at 12 months of age. Mice were group-housed in individually ventilated cages with a constant temperature of 25°C, a 12-hour light/dark cycle and with RM3 expanded chow (Special Diet Services).

For the above mice projects were approved Newcastle University Animal Welfare Ethical Review Board and experiments were conducted in compliance with the UK Home Office (PPL P3FC7C606 or 60/3864).

#### MAO-A transgenic mice

MAO-A mice, on the C57BL6/J background, with cardiac-specific overexpression of MAO-A driven by the α-MHC promoter are previously described(Villeneuve et al., 2013). Mixed sex MAO-A offspring and non-transgenic littermate were used for the experiments at 6 months of age. N-acetyl-cysteine was given at 1.5 g/kg/day between 1 month and 6 months of age. Mice were housed in a pathogen-free facility (B 31 555 010). Experiments were approved by University of Toulouse local ethic committee (**CEEA-122 2015-01) and** conformed to the Guide for the Care and Use of Laboratory Animals published by the Directive 2010/63/EU of the European Parliament.

#### INK-ATTAC mouse model for the clearance of senescent cells

Experimental strategy for making transgenic mice with a senescence-activated promoter coupled to the drug-activatable ATTAC “suicide” transgene and GFP was devised by JLK and TT. INK-ATTAC mice were created and characterized at Mayo through a collaboration among the JLK-TT, NKL, and van Deursen laboratories. Animals were crossed onto a C57BL/6 background (van Deursen laboratory) and bred, genotyped, and aged (JLK-TT laboratory). Mixed sex mice were housed in a pathogen-free facility, at 2–5 mice/cage with 12 h light/12 h dark cycle at 24 °C and ad libitum access to standard mouse diet (Lab Diet 5053, St Louis, MO, USA) and water. AP20187 (10 mg/kg) or vehicle was administered to 27-month-old mice by intraperitoneal injection every 3 days for 2 months.

#### Senolytic treatment of male C57BL/6 mice

Mice at 22 months of age were purchased from Charles River (Charles River Laboratories International, UK). Mice were randomly assigned to a treatment group. ABT263 (navitoclax) or vehicle alone was administered to mice by oral gavage at 50 mg/kg body weight per day (mg/kg/d) for 7 d per cycle for two cycles with a 1-week interval between the cycles. Hearts were collected directly into 50 mM KCl to arrest in diastole.

For all animal studies details of sample sizes is included in the figure legends.

#### Human Tissue Collection and Ethics

Human heart tissue was obtained from male and female patients undergoing open heart surgery for aortic stenosis, with a section of the right atrial appendage being placed in 10% neutral buffered formalin (VWR, 9713.9010) immediately after dissection. Subsequent processing steps for paraffin embedding were the same as for mouse tissue (as described below). All tissue samples were obtained under the clause in the Human Tissue act that enables anonymised samples to be taken without consent in the context of an ethically approved study. This study was approved by the Research Ethics Committee UK, REC reference number: 10/H0908/56.

#### Adult Mouse Cardiomyocyte Isolation and Purification

Hearts from mixed sex CJ57/BL6 mice, were placed on a Langendorff setup. Cell suspensions were obtained by enzymatic digestion and sedimentation. To enrich the CM population, cells were stained at 4°C for 30 min with a cocktail of biotin conjugated antibodies (CD31 (390), CD45 (30-F11), and Sca-1 (D7)) (Biolegend) and the EasySep™ Mouse Biotin Positive Selection Kit, used to remove the labelled non CM cells. Cultured CM purity was assess by immunofluorescent staining against CD31, Sca-1, CD45, and α-actinin. Qiagen RNA extraction kit was used of RNA isolation. CMs were cultured on laminin-coated wells in MEM with Hank’s salts and L-glutamine (SIGMA), 0.1mg/ml BSA, 10mM Butanedione monoxime, ITS Liquid Media Supplement (SIGMA).

#### Culture of H9C2 rat myoblasts

H9C2 rat heart-derived embryonic myocytes (ATCC) were cultured in DMEM (SIGMA), 5% foetal calf serum, penicillin/streptomycin 100 U/ml, 2mM glutamine, at atmospheric conditions (air plus 5% CO2). Cells were passaged prior 70% confluency.

#### Neonatal Rat Ventricular Myocyte (NRVM) Isolation

For neonatal cardiomyocytes, hearts of mixed sex, 2-3 days old Sprague-Dawley rats were dissociated with 0.43mg/mL collagenase type A (Roche) and 0.5mg/mL pancreatine (Sigma), as previously described (Fazal L., Laudette M. et al., Circ Res 2017). Myocyte enrichment was performed by centrifugation through a discontinuous Percoll gradient and resultant suspension of myocytes was plated onto gelatin-coated culture dishes in mix medium containing 80 % DMEM High Glucose and 20 % M199, supplemented with 10 % FBS, 5 % HS, and 1% antibiotics. After plating, cells were cultivated in mix medium with reduced FBS concentration (5%). Plasmid transfections were performed using Lipofectamine 2000 reagent (Life Technologies).

#### Mouse Embryonic Cardiomyocyte Isolation and Culture

Under sterile conditions, hearts were removed from mixed sex E17.5 embryos and then cut into multiple fragments in cardiomyocyte balanced salt buffer (CBSB) (116mM NaCl, 20mM HEPES, 1mM NaH_2_PO_4_, 5.365mM KCl, 831nM MgSO_4_). Heart fragments were then incubated in CBSB with 80U/ml collagenase type II (Worthington) for 5 minutes. Allowing fragments to settle, supernatant was removed and centrifuged at 700×g for 5 minutes. Remaining fragments were then re-suspended in cardiomyocyte enzyme solution (CBSB, 80 U/mL collagenase II and 0.25 mg/mL trypsin) for 30 minutes, with gentle shaking. Fragments were allowed to settle and then supernatant containing dissociated cells was removed and centrifuged at 700×g for 5 minutes, washed in FBS, and stored at 4°C. Remaining fragments were then re-suspended as before into cardiomyocyte enzyme solution. After centrifugation, the supernatant was discarded and pellet was again re-suspended in 4°C FBS and placed on ice. Re-suspension, centrifugation, and collection were then repeated until all fragments had been dissolved (7-10 cycles). All FBS suspensions were pooled together and centrifuged at 700 x g for 5 minutes. The supernatant was then discarded and cells re-suspended in cardiomyocyte growth medium (Dulbecco’s Modified Eagle’s Medium (DMEM), supplemented with 17% Medium 199, 5% FBS, 10% Horse serum (SIGMA, H0146), 100μg.ml^-1^ streptomycin, 100 units.ml^-1^ penicillin, and 2mM l-glutamine). Cells were then seeded into a collagen-coated (1mg.ml^-1^) T75 culture flask (SIGMA) and incubated at 37°C for 2 h. After the incubation the supernatant, containing cardiomyocytes, was collected from the flask and the adherent fibroblasts discarded. Cardiomyocytes were then cultured in Cardiomyocyte Growth Medium.

#### Culture and Differentiation of AC10 Cells

Cells where seeded at a density of 2.5×10^4^/ml on glass cover slips coated with 12.5 μg/ml fibronectin in 0.02% gelatin in DMEM/F-12 with 12.5% FBS. Once confluent, medium was then replaced by mitogen-deficient medium, DMEM/F-12 supplemented with 2% HS and insulin–selenium–transferrin supplement (ITS, Invitrogen), and the cells cultured for a further 2 weeks to allow terminal differentiation.

### Cell Culture, transfections and Treatments

#### Transfection of H9C2 myoblasts

H9C2 cells were transiently transfected with a 53BP1-GFP reporter protein using the plasmid pG-AcGFP-53BP1c or either TRF1-FokI-D450A or TRF1-FokI plasmids using Lipofectamine 2000 (ThermoFisher) at a ratio of 3μL Lipofectamine 2000 to 1μg DNA following the manufacturer’s protocol. Transfected cells detected with an anti-FLAG antibody (SIGMA).

#### Stress-induced senescence of culture cells

For wither H9C2 cells or embryonic CMs senescence was induced by 10Gy X-ray irradiation. In both cell types, we observed senescence after 10 days using SA-□-Gal assay (>70% positive cells) and absence of proliferation marker Ki-67. H9C2 senescent or proliferating cells were treated with ABT-263 (Navitoclax) (AdooQ Bioscience, USA) and viability was assessed using a Tali Image-Based Cytometer (Invitrogen).

#### EdU incorporation assays

Cells were incubated in 10μM EdU in normal growth medium for 24 h. EdU was detected using the Click-iT^®^ EdU Imaging Kit (Invitrogen), as per manufacturer’ protocol.

#### Thoracic Irradiation

A TrueBeam linear accelerator (Varian Medical Systems, Palo Alto) was used for the mouse studies. Anesthetized INK-ATTAC mice (2 months of age) received a single, 20Gy radiation dose delivered to the thoracic region. The radiation beam was collimated to an area encompassing the mouse lungs and the radiation field positions on the mice were verified using kV-CBCT and 2D kV imaging of the animals prior to irradiation. The dose rate at the prescription point was 14.8 Gy/ min, using 89 cm source to the surface distance. Dose was prescribed to midline in the mice and was confirmed using film and ion chamber dosimetry. All mice prophylactically received 100 mg/ml Baytril antibiotic in their drinking water for 3 weeks post-irradiation. One month following irradiation, mice were randomized to AP20187 (10 mg/kg), delivered by intraperitoneal injection (treatments twice weekly) or vehicle groups. Body weight was monitored weekly. Mice were euthanized 6 months post-irradiation exposure using a lethal dose of pentobarbital. Animal experiments involving INK-ATTAC mice were performed under protocols approved by the Mayo Clinic Institutional Animal Care and Use Committee.

#### Live-Cell Imaging

For live-cell time lapse microscopy of 53BP1 reporter fluorescence, H9C2 cells were plated on glass coverslip bottomed dishes (MatTek), and cells were imaged every 10 minutes for 10 h as Z-stacks over 7μm with a 63X objective (NA=1.4), using a Zeiss Spinning Disk confocal microscope with cells incubated at 37°C with humidified 5% CO_2_. AcGFP-53BP1 foci dynamics were analysed using Imaris Software.

#### Immunohistochemistry

Deparaffinization, hydration, and antigen retrieval of formalin-fixed paraffin-embedded heart tissues was performed as previously described. Sections were incubated with M.O.M mouse IgG blocking reagent (90μL blocking reagent: 2.5mL TTBS) for one hour at room temperature. After two times five minute PBS washes, sections were incubated with avidin for 15 minutes, rinsed with PBS, and then incubated with biotin for 15 minutes at room temperature. Primary antibody was then diluted in M.O.M diluents (7500mL TTBS: 600uL protein concentrate) and incubated overnight at 4°C. After 2 × 5 minute PBS washes, sections were incubated in biotinylated-mouse IgG reagent diluted in blocking solution (1:200) for 30 minutes at room temperature. After 2 × 5 minute PBS washes, endogenous peroxidise activity was blocked by incubating sections in 0.9% H2O2 in water for 30 minutes. After 2 × 5 minute PBS washes, sections were incubated in AB-Complex for 30 minutes at room temperature. After 3 × 5 minute PBS washes, sections were then incubated in NovaRed solution for up to 10 minutes, rinsed with water, counterstained with haematoxylin for 2 minutes, washed in PBS and then transferred to ammonia water for 30 seconds. After a 1 minute wash in water, sections were then dehydrated through 70%, 90% and 100% ethanol and then histoclear for 5 minutes each, Sections were then mounted with DPX.

Primary antibodies used: mouse monoclonal anti-4HNE (1:100, MHN; JaICA), mouse monoclonal anti-8-OHdG (1:100, MOG; JaICA), rabbit polyclonal anti-p21 (1:200, ab7960; Abcam).

#### Immunofluorescence

Cells were grown on sterile coverslips, fixed with 2% PFA and permeabilised with PBG-T (PBS, 0.4% Fish-skin Gelatin, 0.5% BSA, 0.5% Triton X-100). Cells were incubated in primary antibody for 1 hour at room temperature (RT), followed by incubation in secondary antibody for 45 minutes. Lists of primary antibodies and secondary antibodies used in Supplementary materials and methods. Methods: anti-γ-H2AX (1:200, #9718; Cell Signalling), anti-53BP1 (1:200, #4937; Cell Signalling), anti-53bp1 (1:200, nb100-305, novus biologicals), anti-α-actinin (1:100, A7811; SIGMA), anti-troponin I (1:200, Catalogue, Company), anti-troponin C (1:200, Abcam ab30807), anti-PCM1 (1:200, ab154142; Abcam), anti-FLAG (1:1000, F1804; SIGMA), anti-Ki67 (1:250 (4μg ml-1), ab15580; Abcam). Secondary antibodies: Alexa Fluor® 488, Alexa Fluor® 594, or Alexa Fluor® 647 (Invitrogen).

#### Microscopy

For fluorescence microscopy a Leica DM5500B wide-field fluorescence microscope and Zeiss AxioObserver Spinning Disk confocal were used. For super-resolution STED microscopy a Leica SP8 confocal (inverted) gSTED 3D super resolution microscope was used. For all quantitative immunohistological analysis studies were performed in a blinded manner.

#### Immuno-FISH and FISH

Immuno-FISH was performed as described (Passos, Saretzki et al., 2007a). Briefly, cells grown on coverslips and immunocytochemistry performed as described above, with either anti-γH2AX (1:200, 9718; Cell Signalling) or anti-53BP1 (1:200, 4937; Cell Signalling). Cells were then fixed with methanol: acetic acid (3:1), dehydrated through an alcohol series, air-dried and incubated with PNA hybridisation mix (70% deionised formamide (SIGMA), 25mM MgCl_2_, 1M Tris pH 7.2, 5% blocking reagent (Roche) containing 2.5 μg/mL Cy-3-labelled telomere specific (CCCTAA) peptide nucleic acid probe (Panagene)), for 10 minutes at 80°C in a then for 2 h at RT in the dark humidifier chamber. Cells were washed with 70% formamide in 2 × SSC for 10 minutes, 2 × SSC for 10 minutes, and PBS for 10 minutes. Cells were then dehydrated an alcohol series, air dried and mounted with ProLong® Gold Antifade Mountant with DAPI (ThermoFisher). For immuno-FISH in formalin-fixed paraffin-embedded tissues, sections were de-waxed and rehydrated through an alcohol series. Antigen retrieval was via 0.1M citrate buffer boiling. Section were incubation in blocked buffer (1:60 Normal Goat Serum in PBS/0.1% BSA) followed by primary antibody at 4°C. Sections were then incubated with a biotinylated secondary antibody (1:200) followed by avidin-DCS (1:500). Sections were post-fixed in 4% PFA for 20 min and FISH was carried out as described above. For centromere FISH (SADS detection) a FAM-labelled, CENPB-specific (centromere) (ATTCGTTGGAAACGGGA) peptide nucleic acid probe (Panagene) was used. Frequency of SADS was assessed by quantification of decondensed/elongated centromeres.

#### RNA *in situ* Hybridization

RNA-*ISH* was performed after RNAscope protocol from Advanced Cell Diagnostics Inc. (ACD). Paraffin sections were deparaffinised with Histoclear, rehydrated in graded ethanol (EtOH), and H_2_O_2_ was applied for 10 min at RT followed by two washes in H_2_O. Sections were placed in hot retrieval reagent and heated for 30 min. After washes in H_2_O and 100% EtOH, sections were air dried. Sections were treated with protease plus for 30 min at 40 °C, washed with H_2_O, and incubated with target probe (p16, eGFP) for 2 h at 40 °C. Afterwards, slides were washed with H_2_O followed by incubation with AMP1 (30 min at 40 °C) and next washed with wash buffer (WB) and AMP2 (15 min at 40 °C), WB and AMP3 (30 min at 40 °C), WB and AMP4 (15 min at 40 °C), WB and AMP5 (30 min at RT) and WB and, finally, AMP6 (15 min at RT). Finally, RNAscope 2.5 HD Reagent kit-RED was used for chromogenic labelling. After counterstaining with haematoxylin, sections were mounted and coverslipped. All analysis was performed in a blinded manner.

#### Crosslinked Chromatin Immunoprecipitation Assay (ChIP)

3 and 30 month old frozen heart tissue (n=3 per age group) was powdered using dry pulverisation cryoprep (Covaris). The powder was then resuspended into 1% formaldehyde solution in PBS and cells cross-linked for 5 minutes at RT, then quenched with glycine buffer at final concentration of 125 mM. The cells were rinsed in cold PBS, and harvested into PBS with protease inhibitors and centrifuged for 5 min, 4°C, 1,000 x g. The pellet was resuspended in ChIP lysis buffer (1% SDS, 10mM EDTA, 50mM Tris-HCl pH8.1) and incubated for 20 min on ice. Lysate was sonicated to generate average size fragment of 200-700bp; cell debris was pelleted by centrifugation for 10 min, 4°C, 8,000 x g. The supernatant contained chromatin, which was quantified using OD_260_ on Nanodrop.

25 μg of chromatin per IP was diluted 1:10 with dilution buffer (1.1% Triton, 1.2mM EDTA, 16.7mM TRIS-HCl pH8.1,167mM NaCl, with protease inhibitors) and precleared by incubating with 25 μl blocked Staph A membranes (Isorbyn) for 15 minutes. Precleared chromatin was then incubated with 5 μg anti-γH2AX (phospho S139) (Abcam, ab2893) and species and isotype matched control ChIP grade IgG (Abcam, ab46540) overnight at 4°C. Immunoprecipitated chromatin was collected with blocked Staph A membranes, which were then sequentially washed with cold buffers; Low Salt Wash Buffer (0.1% SDS, 1% Triton X-100, 2 mM EDTA, 20 mM Tris-HCl pH 8.0, 150 mM NaCl), High Salt Wash Buffer (0.1% SDS, 1% Triton X-100, 2 mM EDTA, 20 mM Tris-HCl pH 8.0, 500 mM NaCl), LiCl Wash Buffer (0.25 M LiCl, 1% NP-40, 1% Sodium Deoxycholate, 1 mM EDTA, 10 mM Tris-HCl pH 8.0) and two washes in TE Buffer (10 mM Tris pH 8.0, 1 mM EDTA). The immunoprecipitated chromatin was eluted in 500 μl Elution buffer (1% SDS,100mM NaHCO_3_), cross-links reversed and DNA obtained by phenol:chloroform extraction and gel purification. Real-time PCR specific for telomeric repeats was performed as described in (50). Briefly, 4 μL of ChIP eluate was added to a final volume of 13 μl quantitative PCR reaction containing 6.5 μL Jumpstart SYBR master mix (Sigma) and 10pmol primer mix. Telomere PCR reaction conditions were: 10 seconds at 95°C followed by 40 cycles of 10 seconds at 95°C, 30 seconds at 55°C, and 30 seconds at 72 °C followed by ABI7500 Fast Real-Time PCR System machine predetermined melt curve analysis (Applied Biosystems). Each PCR reaction was performed in triplicate. All reactions were normalized to the control antibody and fold enrichment above background was calculated using the following equation: (1/(2A)) ×100. The values were then adjusted to the values of total input and expressed as log to the base 2.

#### Senescence-Associated β-Galactosidase Activity Assay

SA-β-Gal activity assay was performed as described previously (Dimri, Lee et al., 1995). For *in vitro* studies cells were grown on sterile coverslips and for *in vivo* studies samples were cryo-embedded and staining performed on 10μm sections. Samples were fixed with 2% paraformaldehyde in PBS for 5 minutes, washed with PBS for 5 minutes and then incubated at 37°C overnight in SA-β-Gal solution containing 150mM sodium chloride, 2mM magnesium chloride, 40mM citric acid, 12mM sodium phosphate dibasic, 5mM potassium ferricyanide, 5mM potassium ferrocyanide, and 1mg ml^-1^ 5-bromo-4-chloro-3-inolyl-β-d-galactosidase (X-Gal) at pH 5.5. For *in vivo* staining and quantifications sections were co-labelled with anti-troponin C and WGA as above. All analysis was performed in a blinded manner.

#### TRAP Assay

Heart samples were snap-frozen in liquid nitrogen immediately after dissection. Using a liquid-nitrogen-cooled pestle and mortar, tissues were ground into a fine powder. Telomerase activity was then determined following the TeloTAGGG Telomerase PCR ELISA kit (Roche).

#### Assessment of cardiomyocyte Hypertrophy

Hypertrophy was assessed by cross-sectional area using established methods as described(Shenje, Andersen et al., 2014). Troponin C was used to identify CMs and membranes labelled with wheat germ agglutinin (WGA) (Alexa Fluor® 647, W32466, Invitrogen). Only CMs in the left ventricle free wall were analysed. To control for tissue orientation only myocytes that were surrounded by capillaries all displaying a cross-sectional orientation were analysed. All analysis was performed in a blinded manner.

#### Evaluation of cardiomyogenesis in mice

To allow retrospective quantification of cumulative proliferation and binucleation, occuring the week following navitoclax treatment, mice were injected (intraperitoneal) with EdU (100 μg/g body weight) once daily every 24 h for 7 consecutive days (**Fig. 7 a**). More than five animals were studied for each treatment group. Edu incorporation was detected using the click-iT™ EdU Imaging Kit (Thermofisher). Sections were also labelled with primary antibody for troponin C and the membrane specific label WGA. A Leica SP5 laser scanning confocal system was used for the analysis. Analysis was carried out in a blinded fashion similar to our previously studies (Richardson et al., 2015). In brief, myocardial tissue was sectioned transversely in 40?μM sections, with sections separated by intervals of 200?μM, allowing representative analysis through-out the ventricle. Over 80 images were taken, in a random fashion, spanning over 8 individual sections per heart. Analysis included the assessment of Edu labelling in >1500 cardiomyocytes per heart. Incorporation of EdU in the cardiomyocyte population was quantified based on the detection of EdU labelling in the nucleus of a troponin C expressing cell. To discriminate proliferation from binucleation (karyokinesis without cytokinesis), the number of EdU labelled nuclei within each Edu^+^ cardiomyocyte was assessed using WGA to identify the cell membrane. The incorporation of EdU into a cell with only one nucleus was considered to be a result of proliferation, when two or more EdU labelled nuclei were found in the same cardiomyocyte, this was considered to have been incorporated during a multinucleation event. Ki67 or Aurora B expressing cardiomyocytes were identified as co-expression of Ki67/Aurora-b and troponin C within a single cell membrane (based on WGA labelling). The total percentage of positively expressing cardiomyocytes was calculated as previously described (Richardson et al., 2015), based on the analysis of >4000 cardiomyocytes per heart. All quantifications were performed blinded using digital image analysis (ImageJ; U.S. National Institutes of Health; http://rsbweb.nih.gov/ij/). All analysis was performed in a blinded manner.

#### MR Imaging and Analysis

Magnetic resonance images were acquired with a horizontal bore 7.0T Varian microimaging system (Varian Inc., Palo Alto, CA, USA) equipped with a 12-cm microimaging gradient insert (40 gauss/cm) and analysed as described previously(Redgrave, Tual-Chalot et al., 2017). Percentage change in wall thickening was calculated from the mean of wall thickness at 4 points in all slices at end systole and end diastole thickness. % change=(EST-EDT)/EDTx100. All analysis was performed in a blinded manner.

#### RNA-Sequencing

Paired-end reads were aligned to the mouse genome (mm9) using a splicing-aware aligner (tophat2). Only unique reads were retained. Reference splice junctions were provided by a reference transcriptome (Ensembl build 67), and novel splicing junctions determined by detecting reads that spanned exons that were not in the reference annotation. True read abundance at each transcript isoform was assessed using HTSeq (Python) before determining differential expression with the tool DESeq2, which models mean-variance dependence within the sample set. Significance was determined using an FDR corrected p-value ≤ 0.05. Alignment statistics for each replicate are included in the supplementary data.

#### Computational Modelling

Two different models were constructed to examine different hypotheses on how TAF accumulate in cells. The models were encoded in the Systems Biology Markup Language (SBML) modelling standard (Hucka, Finney et al., 2003) with the Python SBML shorthand tool (Wilkinson, 2011). Model simulations were carried out in COPASI (Hoops, Sahle et al., 2006) and the results were analysed and plotted in R using ggplot2(Wickham, 2009). We used stochastic simulation (Gillespie Direct Method) in order to be able to account for the variability in the experimental data. The models were deposited in BioModels (Chelliah, Juty et al., 2015) and assigned the identifiers MODEL1608250000 and MODEL1608250001.

#### Cufflinks FPKM Determination

To calculate library normalised read counts for each transcript in every replicate, we used the tool Cufflinks, which returns the proportion of reads per million, which mapped to each gene isoform.

#### Gene Set Enrichment Analysis (GSEA)

We perform a GSEA by creating a ranked list of the DESeq2 results file, where genes are ranked by their log transformed, non-FDR-corrected p-values. This list is then used as an input to the GSEA software, which assesses positive or negative shifts of GO ontology classes in the global distribution of gene expression. This is achieved by the calculation of an enrichment score, which reflects the movement of a particular ontological class to the positive and negative extremes of the distribution of the expression of all genes. Permutation testing then allows for the estimation of significance prior to FDR correction.

#### Heatmaps

Heatmaps were created in R using the *ggplots* package. These heatmaps displaying normalised (row scaled – Z-score) FPKM gene expression across a series of replicates were clustered using a Pearson correlative clustering approach in the “hclust” R package.

#### Principal Component Analysis

Principal component analysis was performed in R using the *prcomp* method.

#### Data deposition

The RNA-seq data from this publication have been submitted to the GEO database (http://www.ncbi.nlm.nih.gov/geo/)

#### Statistical Analysis

We conducted two-tailed *t*-tests, one-way ANOVA, Gehan-Breslow, and Mann-Whitney tests using GraphPad Prism.

## Acknowledgements

This work was funded by BBSRC grants BB/H022384/1 and BB/K017314/1 to JFP, BHF project grant PG/15/85/31744 to GR and JFP, NIH grant AG013925 and funding from the Connor Group and Noaber Foundation to JLK, and grant from Région Midi-Pyrénées, France to JMP, DM, and VDE. CJP and JFP were funded by the MRC-Arthritis Research UK Centre for Integrated research into Musculoskeletal Ageing (CIMA) grant MR/K0063121/1. We thank Simon Tual-Chalot for the expert assistance in magnetic resonance imaging. We thank Prof. Roger Greenberg for kindly providing us with the TRF1-FOKI plasmid. We thank Gabriele Saretzki for expert assistance in TRAP assays, Thomas von Zglinicki for providing mice tissues and Glyn Nelson for assistance in live-cell microscopy. We thank Utz Herbig, Steve Kosak for insights into telomere biology.

## Author Contributions

RA, DM, and AL performed the majority of experiments. JC, JB, HS, MO, DJ, CP, CCM, ED, AW, EF, RBP, KLK, VDE, MJS, CMR,NR, JM, TT performed and evaluated individual experiments; AO, AP, NKLB, JM, JLK, and PDA designed and supervised individual experiments; LG provided materials; JMP, GDR, and JFP designed and supervised the study; JFP, RA, and GDR wrote the manuscript with contributions from all the authors.

## Competing financial interests statement

Patents on INK-ATTAC mice are held by Mayo Clinic and licensed to Unity Biotechnology. J.L.K. and T.T. may gain financially from these patents and licenses. This research has been reviewed by the Mayo Clinic Conflict of Interest Review Board and was conducted in compliance with Mayo Clinic Conflict of Interest policies. The remaining authors declare no competing financial interests.

## Supplementary Data (S1-13)

**Figure S1.**
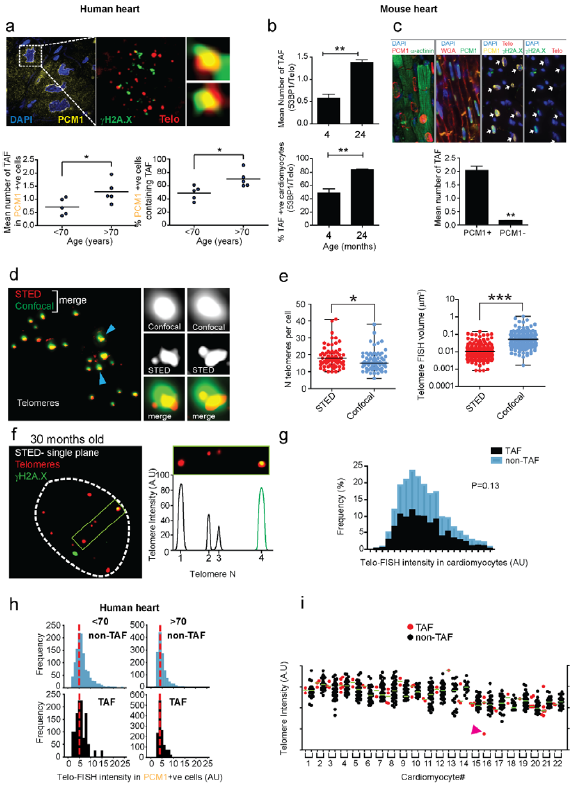
Aged cardiomyocytes in both humans and mice are characterised by dysfunctional telomeres. **(a)** Representative images of γH2A.X immuno-FISH in PCM1-positive human cardiomyocytes (blue – DAPI; yellow – PCM1; red – telo-FISH; green – γH2A.X). Images are z projections of 4.5μM stacks taken with 100X objective. Right panels show co-localisation between telomeres and γH2AX, with taken from single z planes where co-localisation was found. Graphs showing mean number of TAF (left) and mean percentage of TAF positive nuclei (right) in PCM1-positive human cardiomyocytes from 46-65 and 74-82 year old human heart tissue. Data are represented as the mean for individual subjects, with the horizontal line representing group mean. **(b)** Quantification of 53BP1 ImmunoFISH in 4 and 24 months old mice. Graphs representing mean number of TAF and % of cardiomyocytes positive for TAF; n=3 mice per age group; >100 cardiomyocytes were quantified. Asterisk denote a statistical significance at P < 0.05 using two tailed t-test. **(c)** Above: representative images of PCM1 and α-actinin; PCM1 and WGA and γH2A.X, PCM1 immuno-FISH in 30 month old mouse cardiac tissue. Below: Graph representing mean number of TAF in PCM1 positive versus PCM1 negative cardiac cells; n=3 mice; >100 cardiomyocytes were quantified. Asterisk denote a statistical significance at P < 0.05 using two tailed t-test. **(d)** Representative images of conventional confocal (green) versus STED microscopy (red) for detection of telomeres by Q-FISH. Note that STED microscopy is capable of discerning telomere clusters which would otherwise be detected by confocal microscopy as a single signal. **(e)** Comparison between average number of telomere FISH signals detectable per cell (left graph) and mean telomere volume (right graph) in mouse cardiomyocytes, detected by either standard confocal or STED microscopy. Data are represented as the mean for each measurement, with the horizontal line representing the group mean. **(f)** Representative image of immuno-FISH using STED microscopy for γH2A.X and telomeres in cardiomyocytes from a 30 month old mouse. Graph on the right side shows 2 telomere FISH signals of similar intensities (one showing co-localisation with γH2A.X and the other not). **(g)** Histograms representing Q-FISH analysis by 3D STED microscopy comparing individual telomere length either co-localising (TAF) or not co-localising (non-TAF) with γH2AX foci in mouse cardiomyocytes from n=3 mice (aged 30 months of age). 300 cardiomyocytes (detected by troponin C and WGA) were analysed per mouse. Mann-Whitney test reveals no statistical significance between TAF and non TAF P=0.13. **(h)** Graph showing values for telomere FISH intensities either co-localising (TAF) or not co-localising (non-TAF) with γH2AX foci in 22 individual cardiomyocytes chosen randomly. Purple arrow indicates 1 telomere FISH signal which is lower in intensity than the median. **(i)** Histograms displaying telomere intensity for telomeres co-localising (bottom) or not co-localising (top) with γH2A.X DDR foci for PCM1 positive cardiomyocytes obtained from 46-65 (left) and (74-82) year old subjects. Dotted lines represent median intensity. Mann-Whitney tests show no significant difference in telomere intensity between TAF and non-TAF in either, 46-65 or 74-82 year old subjects (P>0.05). More than 100 cardiomyocytes were quantified per subject.

**Figure S2.**
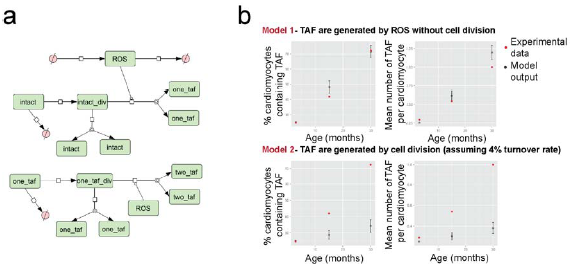
Stochastic mathematical models representing a cardiomyocyte population throughout 27 months of a mouse’s lifespan. **a)** Network diagram of **Model 2**,-TAF are generated by cell division. Reactions involving turnover of cells with two, three or four TAF are similar to those for cells with one TAF and are omitted for clarity. Network diagram created in CellDesigner using Systems Biology Graphical Notation. **b)** Middle and bottom panel: model output showing mean values of 100 stochastic simulations, (error bars show ± one s.d. from the mean).

**Figure S3.**
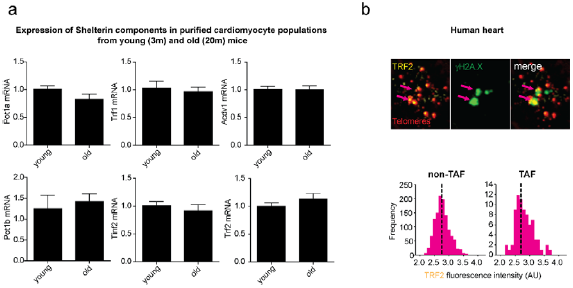
Expression of shelterin components does not change with age in cardiomyocytes. **(a)** mRNA expression of shelterin components: Pot1a, Pot1b, Trf1, Trf2, Tinkf2, and Acdv1 in purified cardiomyocyte populations from young (3m) and old (20m) wild-type mice. Data are mean ± S.E.M of n=5 mice per group. Two-tailed t-test shows no significant difference in expression (P>0.05). **(b)** Representative image of TRF2 immuno-FISH in PCM1-positive human cardiomyocytes (blue – DAPI; yellow – TRF2; red – telo-FISH; green – γH2A.X). Images are z projections of 4.5μM stacks taken with 100X objective. (below) Histograms displaying TRF2 fluorescence intensity for TRF2 foci co-localising with telomeres also co-localising with γH2A.X (right) or telomeres not co-localising with γH2A.X (left) in human cardiomyocytes. Red dotted lines represent median intensity. Mann-Whitney tests show no significant difference in TRF2 intensity between TRF2 abundance at TAF and non-TAF (P>0.05).

**Figure S4.**
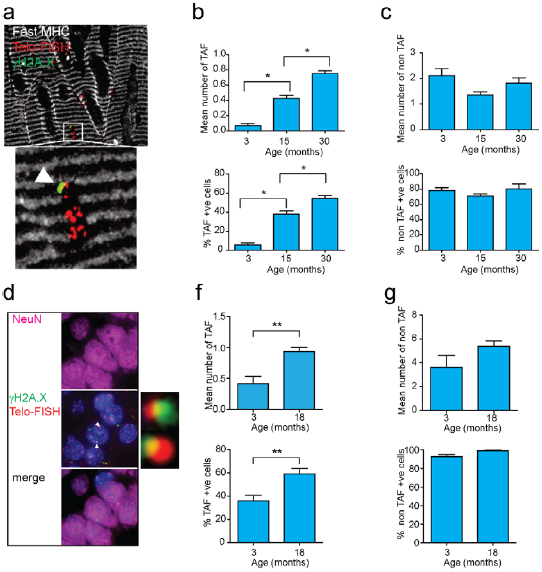
Telomere-dysfunction increases with age in Post-mitotic cells such as muscle fibres and neurones. **(a)** Representative image of Immuno-FISH for γH2A.X, telomeres and fast MHC (marker of type 2 muscle fibres). White arrow indicates co-localisation between γH2A.X and telomeres; **(b)** Mean number of TAF (top) and % of myocytes positive for TAF (bottom); **(c)** Mean number of non-TAF (top) and % of myocytes containing non-TAF (bottom) in 3,15, and 30 month old mice. Data are mean ± SEM of n=4 per age group. More than 70 myocytes were quantified per age group. Statistical analysis performed using One Way ANOVA; * P<0.05; **(d)** representative image of Immuno-FISH for γH2A.X, telomeres and NeuN (neuronal marker) in mouse hippocampus. White arrows indicate co-localisation between γH2A.X and telomeres; **(f)** Mean number of TAF (top) and % of neurons positive for TAF (bottom); **(g)** Mean number of non-TAF (top) and % of neurons containing non-TAF (bottom) in 3 and 18 month old mice. Data are mean ± SEM of n=5-7 per age group. More than 100 neurons were quantified per age group. Statistical analysis performed using two-tailed t-test; * P<0.05.

**Figure S5.**
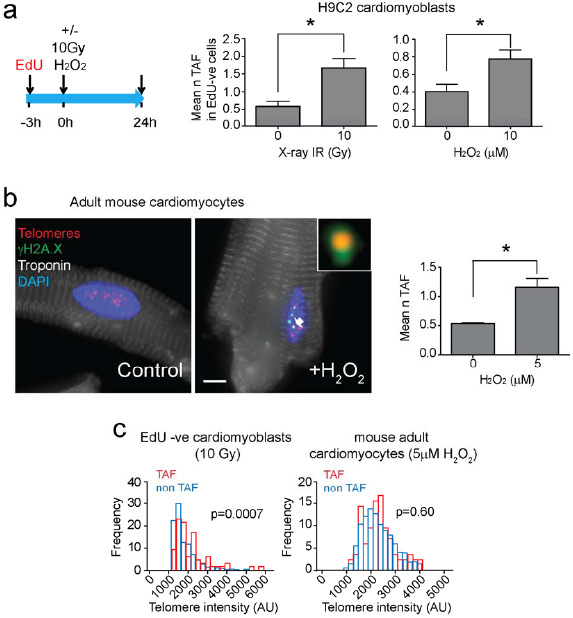
Telomere damage occurs irrespectively of cell division. **a)** Schematic illustration depicting H9C2 cells were incubated in 10μM EdU in normal growth medium for 3 hours, followed by either 10Gy X-irradiation or incubation with 10μM H_2_O_2_ and cultured for a further 24 hours in the presence of 10μM EdU in normal growth medium. Mean number of TAF in EdU positive cells following X-irradiation (left graph) or H_2_O_2_ (right graph). Data are mean ± SEM of *n*=3 independent experiments. 50 cells were analysed per experiment. Statistical analysis performed using two-tailed t test; * P<0.05. **b)** (left) Representative micrographs of Immuno-FISH for γH2A.X, telomeres and Troponin C in control and H_2_O_2_ treated adult mice cardiomyocytes. Arrow indicates co-localisation between telomeres and γH2A.X. Scale bar corresponds to 10μm. (right) Mean number of TAF in isolated mouse adult cardiomyocytes 24hr after treatment with 5μM H_2_O_2_. Data are mean ± SEM of adult cardiomyocytes isolated from 3 month-old C57BL/6 wild-type mice. Statistical analysis performed using two-tailed t test; * P<0.05. **c)** Histograms representing Q-FISH analysis comparing individual telomere intensities of telomeres either co-localising (TAF) or not co-localising (non-TAF) with γH2AX DDR foci in 10Gy irradiated EdU-negative H9C2 cardiomyoblasts (left graph) or 5μM H_2_O_2_ treated mouse adult cardiomyocytes from 3 month-old C57BL/6 wild-type mice (n=3 mice). More than 100 cells were analysed per experiment. Statistical analysis performed using Mann Whitney test.

**Figure S6.**
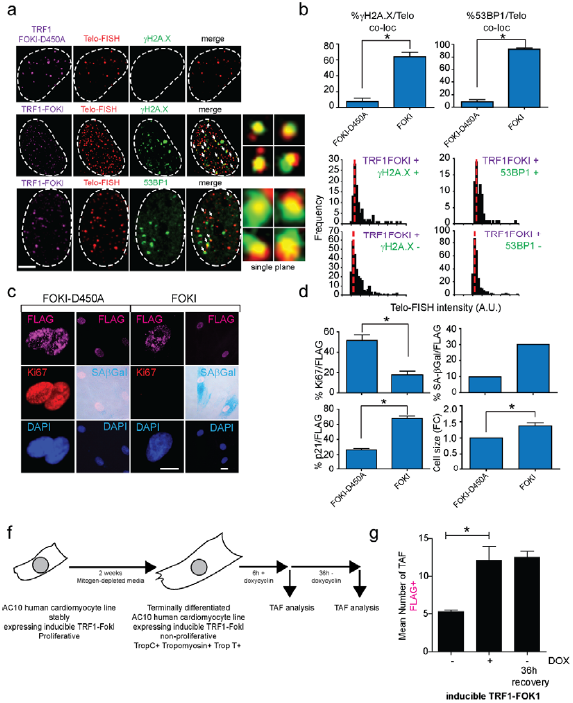
TRF1-FokI Fusion Protein Induces Telomere Specific Double Strand Breaks, senescence and hypertrophy in H9C2 cardiomyoblasts. **(a)** Representative images of H9C2 cardiomyoblasts 4 days following transfection with a FLAG-tagged TRF1-FokI-D450A (top row) or TRF1-FokI (middle and bottom row) fusion protein (cell treatments the same for all subsequent panels in figure) (purple – FLAG; red – telo-FISH; green – γH2A.X/53BP1). Images are z-projections of 0.1μM stacks taken with 100X objective. White arrows indicate co-localisation between telomeres and γH2A.X/53BP1, with co-localising foci amplified in the right panels (taken from single z-planes where co-localisation was found). Scale bar represents 25μm. **b)** Percentage of γH2A.X (left) or 53BP1 (right) foci co-localising with telomeres. Data are mean ± SEM of n=3 independent experiments. More than 50 cells were analysed per condition. Statistical analysis was performed by two-tailed t-test * P<0.05. **c)** Histograms displaying telomere intensity for telomeres co-localising (bottom) or not co-localising (top) with γH2AX (left) or 53BP1 (right) DDR foci. Red dotted lines represent median. Statistical analysis performed using two tailed t test; * P<0.05; More than 50 cells were analysed per condition. Mann-Whitney tests show no significant difference in telomere intensity between TAF and non-TAF, with either γH2A.X (left) or 53BP1 (right) DDR foci. **d)** Representative images of detection of senescent markers (purple – FLAG; red – Ki-67; light blue – DAPI; darker cytoplasmic blue – SA-β-Gal). Scale bar represents 100μm. **e)** Mean percentage of FLAG-positive cells positive for Ki-67, p21, SA-β-Gal activity and cell size. Data are mean ± SEM of n=3 independent experiments. More than 100 cells were quantified per condition. For SA-β-Gal data are presented as mean of >100 cells representative of 1 experiment. Two additional independent experiments confirmed these findings (not shown). Statistical analysis performed using two tailed t test; * P<0.05. **f)** Scheme depicting experimental setting: AC10 human cardiomyocyte cell line stably expressing an inducible TRF1-FokI was cultured for 2 weeks in mitogen-depleted media which led to terminal differentiation and loss of proliferation. Then, cells were treated for 6 hours with doxycycline (DOX), after which, DOX was removed and cells were allowed to recover for 36hours. **g)** Mean number of TAF following induction of TRF1-FokI and recovery. Data are mean ± SEM of n=4 independent experiments.

**Figure S7.**
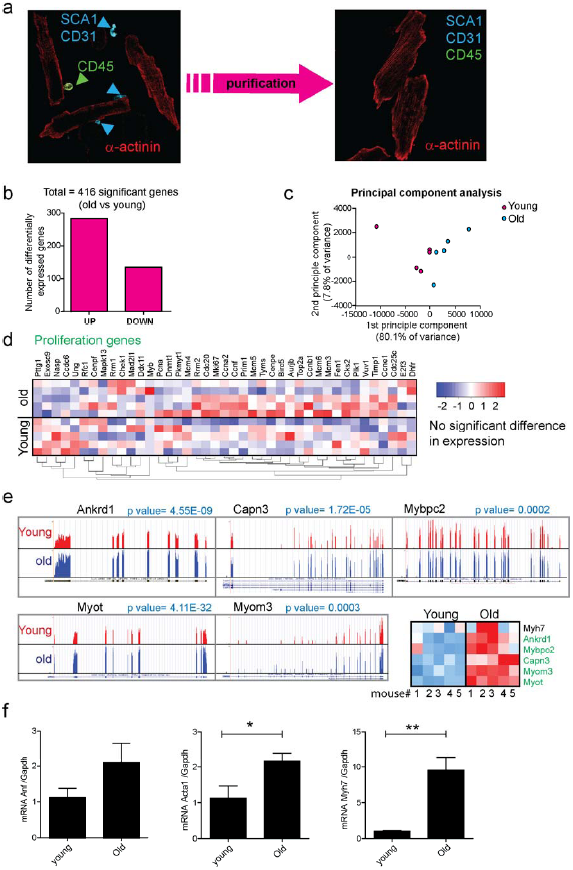
Aged cardiomyocytes have no alterations in proliferation gene expression but show increased expression of genes associated with hypertrophy. **(a)** Representative images of cardiomyocytes isolations before (left) and after (right) purification to remove SCA1-, CD31- and CD45-positive cells. **(b)** Counts of differentially expressed genes calculated using DESeq2 at %5 FDR. **(c)** Principal component analysis (PCA) of FPKM expression for young (red) and old (blue) cardiomyocytes. Components one and two account for 80.1% and 7.8% of the total variance respectively. **(d)** Correlation clustered heatmap of a curated list of known proliferation genes in young and old cardiomyocytes. The colour intensity represents column Z-score, with red indicating high and blue low expression. Note that there is no enrichment for differential expression in this subset of pro-proliferation genes. **(e)** Trace plots and heatmap of the relative expression of Ankrd1, Capn3, Mybcp2, Myot, Myom3 and their associated FDR corrected q-values derived from DESeq2. **(f)** Confirmatory RT-PCRs showing changes in cardiac hypertrophy genes Acta 1 and Myh7, but not Anf. Data were normalised to Gapdh. Data are mean ±S.E.M of n=4-5 mice per age group. Two-tailed t-test was used. **P<0.01; * P<0.05.

**Figure S8.**
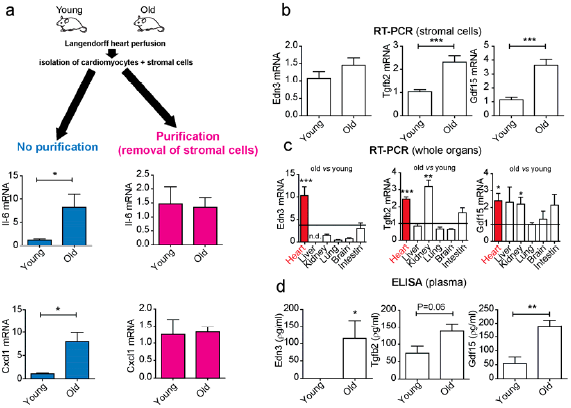
Senescent cardiomyocytes have a non-canonical SASP phenotype. **(a)** Stromal cells isolated as part of the *Langendorff* heart perfusion contribute to the previously reported increased SASP in aged cardiomyocytes: (left) Using the traditional cardiomyocyte isolation method, significant increases in mRNA expression of SASP components Il-6 and Cxcl1 are observed with age; (right) Following removal of stromal cells, no significant differences could be found. Data are mean ±S.E.M of n=4-6 mice per group; **(b)** Stromal cells (no cardiomyocytes) show age-dependent increased expression of Tgfb2 and Gdf15 but not Edn3. Data are mean ±S.E.M of n=8 mice per group. **(c)** Only Edn3 shows an age-dependent increase in expression in the heart. Tgfb2 and GDF15 increase significantly both in heart and kidney. Data are mean ±S.E.M of n=4-7 mice per group. **(d)** Edn3, Tgfb2, and Gdf15 increase with age at the protein level in plasma. Statistical analysis: for multiple comparisons One-way ANOVA was used, otherwise two-tailed t-test. ***P<0.0001; **P<0.01; * P<0.05.

**Figure S9.**
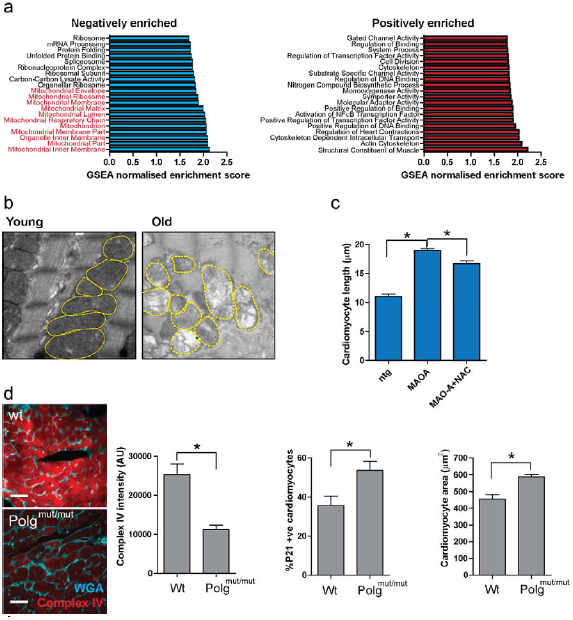
Mitochondrial dysfunction induces senescence and hypertrophy in cardiomyocytes. **(a)** Normalised GSEA enrichment scores for the top 20 positively and negatively enriched GO ontologies. Those negatively enriched ontologies associated with mitochondrial structure and function are highlighted in red. **(b)** Transmission electron microscopy to detect mitochondrial ultrastructural defects in young (3 months) and old (20 months) mice. **(c)** Cardiomyocyte length analysis on WT, MAO-A or MAO-A mice with or without drinking water supplemented with 1.5g/kg/day NAC from the age of 4 to 24 weeks. Data are mean ± S.E.M of n=5-10 mice per group. **(d)** Representative images of mitochondrial complex IV (red) and WGA (blue) in 12 month old WT and Polg^mut^/^mut^ mice. Graphs represent complex IV intensity (left), p21 positivity by immunohistochemistry and mean cardiomyocyte area (right). Data are mean ±S.E.M of n=3-5 mice per group. More than 100 cardiomyocytes were quantified per group. Asterisks denote a statistical significance at P < 0.05 using two-tailed t-test.

**Figure S10.**
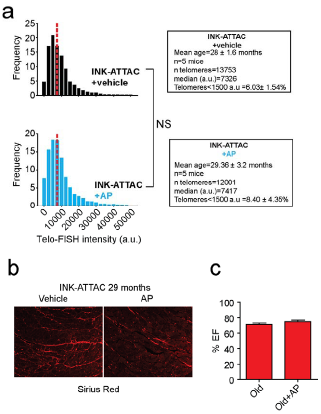
Clearance of senescent cells does not impact on cardiomyocyte telomere length or cardiac function (%EF) but reduces fibrosis. **a)** Histograms showing distribution of individual telomere intensities measured by Q-FISH comparing TAF and non-TAF in INK-ATTAC mice (28-29m old) treated with vehicle or AP20187. >150 cardiomyocytes were analysed per mouse. **b)** Representative image of Sirius Red staining in old INK-ATTAC mice treated AP20187 (or Vehicle) at 29 months of age; **c)** Ejection fraction in old INK-ATTAC treated with or without AP. Data are mean± S.E.M of 11-12 mice per group.

**Figure S11.**
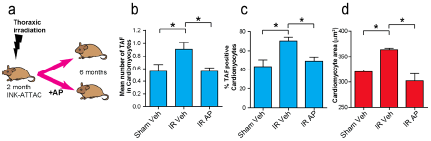
Genetic clearance of p16Ink4a positive cells rescues telomere dysfunction and cardiac hypertrophy after thoracic irradiation. **(a)** Schematic depicting experimental design for figures e-g: 2 month old INK-ATTAC mice underwent thoracic X-irradiation, were treated with AP20187 (or Vehicle) and then sacrificed at 6 months of age; **(b)** Mean number of TAF; **c)** percentage of TAF-positive cardiomyocytes and **(d)** mean cardiomyocyte area μm^2^ in sham irradiated, and irradiated mice with or without AP treatment. Data are mean ± SEM of *n*=6-10 per age group. More than 100 cardiomyocytes were quantified per group. Asterisks denote P < 0.05 using one-way ANOVA;

**Figure S12.**
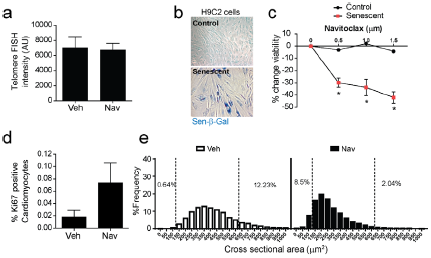
Pharmacological clearance of senescent cells using navitolax does not impact on telomere length but stimulates cardiomyocyte regeneration. **(a)** Mean telomere FISH intensity in 23 month old mice with or without navitoclax treatement. Data are mean ± SEM of *n*=3 per treatment group; **b)** H9C2 cardiomyoblasts were induced to senescence (representative Sen-β-Gal image) via exposure to 10 Gy X-ray radiation; **c)** Navitoclax treatment significantly reduced cell viability in senescent cardiomyoblasts in a dose dependent manner. Navitoclax had no effect on the viability of non-senescent cardiomyoblasts. Data are mean ± SEM n=3 for each treatment group and dosage. Statistical analysis via two-way ANOVA. **d)** Quantification of Ki67 positive cardiomyocytes in vehicle and navitoclax treated animals. Data are mean±S.E.M of n=3 per treatment group; **e)** Distribution of cardiomyocyte cross-sectional area in 23 month old mice treated or not with navitoclax. Data shows that most cardiomyocytes with areas above 650μm^2^ are not detected following navitoclax and smaller cardiomyocytes with areas below 100 μm^2^ emerge. 7 animals per group were analysed.

**Figure S13-.**
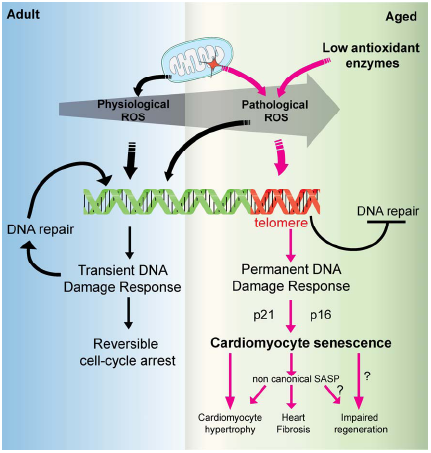
Model to explain cardiomyocyte senescence. In young adult mice, a continuous cycle of “physiological” low level ROS maintains a transient DDR which is a contributor to the low myocardial turnover. During ageing, mitochondrial dysfunction and low expression of antioxidant enzymes, induces “pathological” high ROS which randomly induces telomere-dysfunction that results in a permanent DNA Damage Response sufficient to robustly activate p21 and p16 senescence pathways. Cardiomyocyte senescence (CS) is a driver of myocardial hypertrophy and is characterised by a distinct SASP which contributes to fibrosis and hypertrophy. CS may contribute to impaired regeneration; however the latter hypothesis remains to be tested experimentally.

